# Persistent bacterial coinfection of a COVID-19 patient caused by a genetically adapted *Pseudomonas aeruginosa* chronic colonizer

**DOI:** 10.1101/2020.08.05.238998

**Authors:** Jiuxin Qu, Zhao Cai, Yumei Liu, XiangKe Duan, Shuhong Han, Yuao Zhu, Zhaofang Jiang, Yingdan Zhang, Chao Zhuo, Yang Liu, Yingxia Liu, Lei Liu, Liang Yang

**Affiliations:** Department of Clinical Laboratory, Shenzhen Third People’s Hospital, Southern University of Science and Technology, National Clinical Research Center for Infectious Diseases, 518000, Shenzhen, Guangdong, China; School of Medicine, Southern University of Science and Technology, Shenzhen, 518055, China; The state key laboratory of respiratory diseases, the first affiliated hospital of Guangzhou Medical University, 510120, Guangzhou, Guangdong, China; Medical Research Center, Southern University of Science and Technology Hospital, Shenzhen, Guangdong Province, China, 518055, School of Medicine, Southern University of Science and Technology, Shenzhen, Guangdong Province, China, 518055

**Author notes:** These authors contributed equally to this study (Jiuxin Qu, Zhao Cai). Corresponding authors: Lei Liu, Liang Yang.

**Keywords:** *Pseudomonas aeruginosa*, biofilm, quorum sensing, COVID-19, coinfection

## Abstract

This study characterized a genetically adapted *Pseudomonas aeruginosa* small colony variant isolated from a COVID-19 patient who suffered persistent bacterial coinfection and eventually recovered from critical illness. Specification and modification of the isolates discovered at genomic and transcriptomic levels with aligned phenotypic observations indicated that these isolates formed excessive biofilm with elevated quorum sensing systems.

## Introduction

The characteristics, pathogenicity, epidemiology and treatments of SARS-CoV-2 viral infection has been extensively studied since the onset of the 2019 novel coronavirus (COVID-19) pandemic. [1–3] COVID-19 infection leads to huge impact on the immunity of patients’ respiratory tracks and causes multiple organ damages.[4, 5] Viral infection in the respiratory tract is often accompanied with co-infecting bacterial pathogens. Viral invasion to the respiratory system alters functions and composition of respiratory microbiota which often leads to the onset of secondary bacterial infection. Bacterial coinfection significantly increases the mortality rate of patients while *Pseudomonas aeruginosa* was identified as one of most common co-infecting bacteria during viral infection in respiratory system such as H1N1 pneumonia.[6–8] *P. aeruginosa* is well known to adapt to the respiratory environments by genetic modification to reduce virulence, increase antimicrobial resistance and enhance biofilm formation in immunocompromised individuals with chronic diseases such as cystic fibrosis.[9, 10] However, survival of *P. aeruginosa* in the SARS-CoV-2 infected respiratory tract and its impact on the patients still remain unclear. To investigate the adaptation of *P. aeruginosa* during coinfection with SARS-CoV-2 virus, we characterized two *P. aeruginosa* small-colony variants (SCVs) isolated from sputum samples of a COVID-19 patient who was eventually recovered from critical illness.

## Methods

### Sample collection

Two isolates of *P. aeruginosa* were collected from sputum samples of one COVID-19 patient with 10 days interval (Isolate 1 on 12 February 2020 and Isolate 2 on 22 February 2020) during routine clinical tests. Bacterial antimicrobial susceptibility of *P.aeruginosa* with different colony morphologies, including ceftazidime, piperacillin, cefoperazone/sulbactam, imipenem, aztreonam, and levofloxacin, were performed using the Kirby-Bauer disc- diffusion method. The protocol of K-B method and interpretations of results were based on the Clinical and Laboratory Standards Institute [11]. Susceptibility was listed in Table S8.

### Genomic and Transcriptomic Sequencing Analysis

Genomes of both *P. aeruginosa* isolates were extracted and sequenced by Illumina HiSeq X platform and genome of Isolate 1(Iso1) was sequenced by PacBio Sequel II platform. RNA of two isolates and the *P. aeruginosa* reference PAO1 strain were collected in duplicate and sequenced by Illumina NovaSeq platform. Resequencing analysis was performed to identify Single Nucleotide Polymorphism (SNP) in isolates. PacBio sequence reads was assembled to full genome of Iso 1. Multilocus sequence typing (MLST) was performed to classify the isolates while antimicrobial resistance genes (ARGs) of the isolates were identified. Phylogenetic tree and genomic islands were predicted to trace the origin and identify the specific genomic regions of the isolates. RNA-seq analysis and GO enrichment were performed to investigate the differential gene expression and functional enrichment of the isolates.

### Phenotypic tests

All strains and plasmids used are listed in Table S1. Overnight cultures of the isolates and PAO1 were diluted and incubated for biofilm formation assay. Biofilm biomass was quantified using crystal violet (CV) staining. Swimming and swarming motility of these strains were tested. Quorum sensing (QS) activities of the isolates, PAO1 and PAO1*ΔlasIΔrhlI* mutant (QS negative control strain) was assessed using sterile supernatants of these strains and *PAO1ΔlasIΔrhlI* double mutant harboring either *lasB-gfp* or *rhlA-gfp* reporter. Levels of cyclic-di-GMP in the isolates and PAO1 were measured by monitoring GFP/OD level of transformants of these strains carrying *cdrA-gfp* reporter. All methods are fully described in supplementary documents.

## Results and Discussion

### Patient information

The patient was admitted to our hospital with confirmed COVID-19 infection on 23 January 2020 and hospitalized for 45 days with course of disease of 47 days. The patient received combined antiviral, antimicrobial and anti-inflammatory treatments during hospitalization with immunity enhancing medication and traditional Chinese medicines. (Table S2) Counts of white blood cells (WBC), neutropihls (N), lymphocytes(L) and level of Interleukin-6 (IL-6) were recorded as immune response indicators. (Table S2) The patient turned into critical illness from day 4 to day 32. The first immune response peak was seen at day 7 of hospitalization.(Figure 1S) After receiving treatments, immune response stablized for about 2 weeks and elevated again from day 21 to day 31 with a milder peak.(Figure 1S) The onset of second immune response coincided with the isolation time of *P. aeruginosa* from sputum samples. *P. aeruginosa* persisted in the patient’s respiratory system and could not be eliminated by antimicrobial treatments during the period of critical illness. This suggests that *P. aeruginosa* may play an important role in SARS-CoV-2 coinfection during critical illness and lead to secondary infection. It is a possible cause of the second peak of immune response observed. Thus, two *P. aeruginosa* isolates were collected on day 21 (Iso 1) and day 31 (Iso 2) for further analysis to understanding their genetic and phenotypic characteristics leading to such observation which may contribute to the prognosis of disease development and treatment scheme decision.

### Genotypic and phenotypic analysis reveal that the isolates are SCVs with excessive biofilm formation and elevated quorum sensing

The morphology of the two isolates revealed that they are SCVs of *P. aeruginosa* with overproduction of exopolysaccharides. (Figure 1E) MLST analysis showed that both isolates are *P. aeruginosa* ST 1445, indicating they are the same strain with different evolving times. The complete genome of Iso 1 were assembled and closed with a total length of 6,552,237 bp, average 66.21 % GC and 6239 genes predicted.

**Figure 1.**
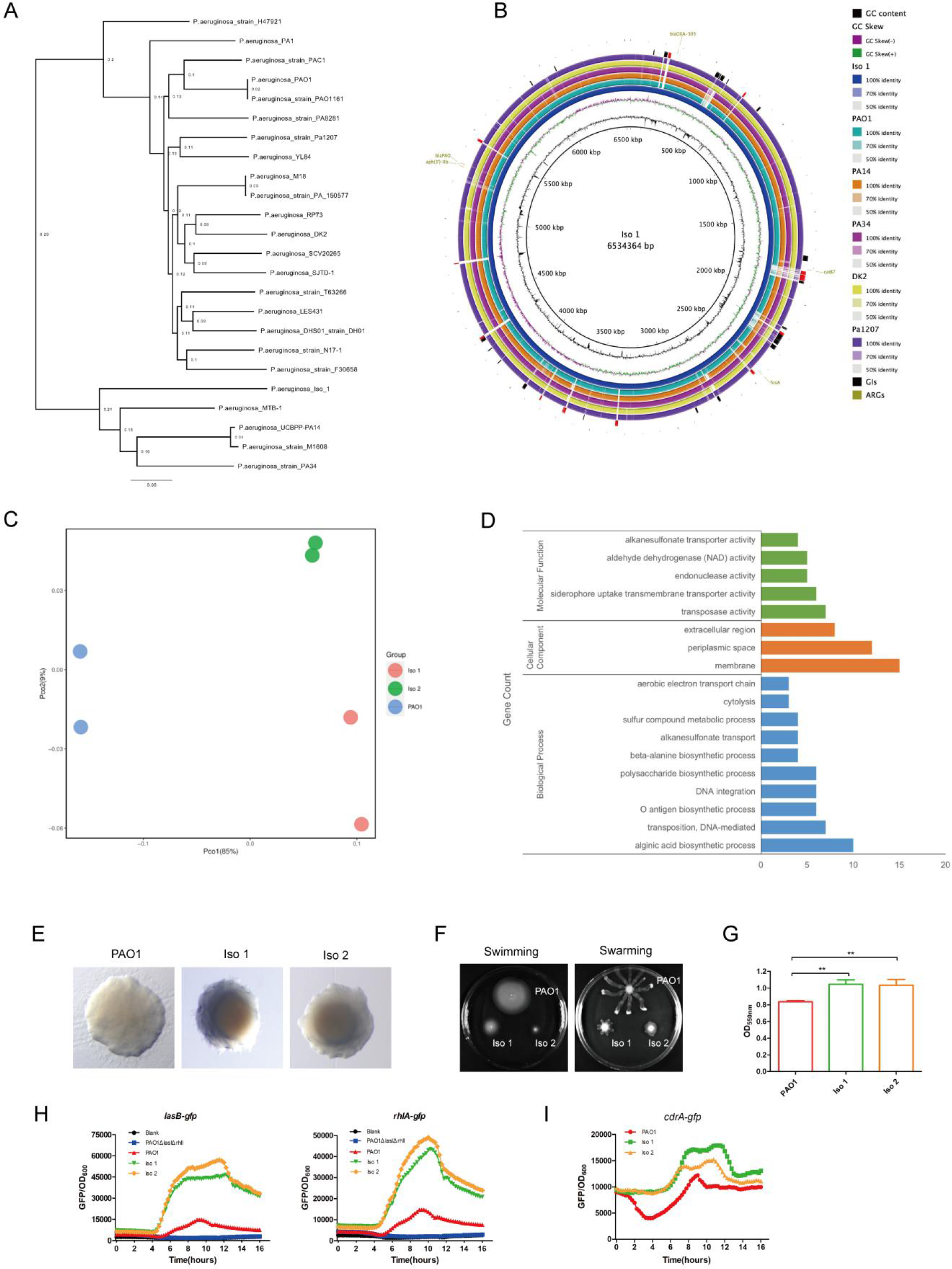
**(1A)** Phylogenetic tree constructed using whole genomes of Iso 1 and 23 other *P. aeruginosa* laboratory, clinical and environmental strains selected from NCBI database; **(1B)** Circular plot. From the innermost, Circle 1: GC content; Circle 2: GC Skew; Circle 3: Iso 1; Circle 4: PAO1(lab reference strain); Circle 5: PA14(virulent clinical isolate); Circle 6: PA34 (strain isolated from Keratitis patient); Circle 7: DK2 (Strain isolated from Cystic Fibrosis patient); Circle 8: Pa1207(strain isolated from Bacteremia patient); Circle 9: Genomic Islands predicted with specific GIs of Iso 1 highlighted in red; Circle 10: Antimicrobial resistance genes; **(1C)** PCoA plot of PAO1, Iso 1 and Iso 2 by Bray Curtis dissimilarity based on normalized read counts of RNA-seq analysis; **(1D)** Gene Orthology Enrichment based on the significantly regulated genes (absolute fold change ≧4, adjusted p-value<0.05) in Iso 2 comparing to that of PAO1. **(1E)** morphology of PAO1, Iso 1and Iso 2; **(1F)** Swimming and Swarming motility of PAO1, Iso 1 and Iso 2; **(1G)** Biofilm formation quantified by CV staining (p-value <0.05); **(1H)** *las* and *rhl* quorum sensing expression measured by *lasB-gfp* and *rhlA-gfp* reporters; **(1I)** Cyclic-di-GMP level measured by *cdrA-gfp* reporter.

Phylogenetic tree was constructed by comparing Iso 1 genome to 23 other *P. aeruginosa* genomes selected from NCBI Genbank to trace its evolutionary origin. (Table S3) Iso 1 branched out individually with a closer distance to the virulent clinical strain PA14 and further from PAO1 and cystic fibrosis strain DK2. (Figure 1A) Comparison of Iso 1 genome with selected genomes of PAO1, PA14, PA34, DK2 and Pa1207 showed that there are specific regions present only on Iso 1 genome. (Figure 1B) Results of genomic islands prediction revealed that these specific GIs (Figure 1B, highlighted in red) on Iso 1 genome including genes involved in transposition protein synthesis, DNA restriction-modification, virulence and toxin transport, and transcriptional regulation, indicating that these specific GIs (Table S4) are important for Iso 1 to survive, persist and invade in the respiratory tract during coinfection. The isolates carry ARGs including *aph(3’)-IIb, bla_OXA-395_, bla_PAO_, fosA* and *catB7* against aminoglycoside, beta-lactam, fosfomycin and phenicol drugs. (Figure 1B, circle 10) These ARGs are crucial for the resistance of the isolates to antimicrobial treatments. However, none of these genes was found locating in the predicted GIs.

To understand the genetic modification of the *P. aeruginosa* isolates evolved over time during SARS-CoV-2 coinfection, SNPs were identified by comparing the genomic sequences of Iso 2 to Iso 1.(Table S5) A total of 118 SNPs were identified from Iso 2 including 22 non-synonymous and 36 synonymous substitutions. Most of the SNPs were found from genes involved in pyoverdine biosynthesis, secretion systems and peptide synthase. Ratio of non-synonymous/synonymous substitutions (<1) indicated a negative selective pressure to the isolates during evolution.

To further understand the adaptation mechanisms of these two *P. aeruginosa* isolates, we compared the transcriptomes of Iso 1, Iso 2 and the reference PAO1 strain. The clear separation of the isolates from PAO1 along PC1 (85%) on PCoA plot implied the significant differences in their transcriptomic profiles.(Figure 1C) The proximity between the clusters of two isolates along both PC1 (85%) and PC2 (9%) implied their transcriptional similarity. Results of differential gene expression (DGE) of the isolates demonstrated that only 39 genes were significantly downregulated (fold change ≦– 4 and adjusted p-value < 0.05) in Iso 2 comparing to Iso 1 with mostly low mean counts. (Table S6) This showed that the isolates survived and proliferated stably during coinfection in COVID-19 patient’s respiratory system. Notably, two downregulated genes with high mean counts are involved in multidrug efflux system(*mexC* and *mexD*), indicating a lower capacity of antimicrobial resistance of Iso 2 comparing to Iso 1. Such change may contribute to the eventual clearance of *P. aeruginosa* by antimicrobial treatments. SNPs identified from Iso 2 had no significant effect on transcriptional expression. DGE of PAO1 was comparing with Iso2 only since there was only minimal difference between Iso 1 and Iso 2. 456 genes were significantly regulated in Iso 2 (absolute fold change ≧ 4 and adjusted p-value < 0.05, Table S7) while Gene Ontology(GO) enrichment were illustrated in Figure 1D. In Iso 2, *cdrA*(PA4625) was upregulated for 4.27 folds indicating an increased cyclic-di-GMP biosynthesis.[12] (Table 1, Figure 1I) This increase in cyclic-di-GMP was most probably due to downregulation of *arr* gene which encodes for a phosphodiesterase catalyzing degradation of cyclic-di-GMP.[13] Moreover, PA2771 encoding a diguanylate cyclase for cyclic-di-GMP biosynthesis also downregulated significantly probably due to its self-inhibition induced upon binding with cyclic-di-GMP.[14] Cyclic-di-GMP promotes biofilm formation through various mechanism including increasing exopolysacchairdes formation and motility reduction. Iso 1 and Iso 2 formed excessive biofilm comparing to PAO1 as observed from phenotypic test.(Figure 1E&G) GO enrichment showed that biosynthesis processes of alginate and lipopolysaccharides (LPS) were highly enriched. (Figure 1D) Genes involved in biosynthesis of alginate (*alg*) and Pel (*pel*) was highly upregulated while those involved in LPS formation (*wbp, wzx/y/z*) were greatly downregulated. Previous studies had proved that alginate, Psl and Pel share the same precursors, mannose-1-phosphate, with LPS.[15] Thus, the increase in the exopolysaccharide of the isolates was due to the biosynthetic flux towards alginate and Pel leading to a reduction in LPS biosynthesis. As LPS is a virulence factor inducing host immune response, the immune response induced in the patient was probably not due to LPS expression. [16] Genes involved in flagellar formation were downregulated in Iso 2 leading to lower swimming and swarmming motility which further contributed to biofilm formation.(Table 1, Figure 1F) An notable observation from the isolates was the upregulation of *las* and *rhl* QS systems.(Figure 1H) Such observation was opposite from previous discovery which showed SCVs with more biofilm formation possessed suppressed QS systems to escape from host immune clearance.[17, 18] From DGE results, expression of *lasB* and *rhlAB* genes in Iso 2 were slightly upregulated for 1.62, 2.74 and 3.25 folds, respectively, which aligned with phenotypic observation. As QS is related to virulence, such observation possibly inferred that the isolates would have attacked the host tissue during critical illness and thus induced host immune responses instead of escaping from immune clearance by suppressing QS. We further found that erythromycin was effective in reducing such elevation of QS. (Figure S2)

**Table 1.** Differential expression of genes involved in cyclic-di-GMP biosynthesis, biofilm formation and motility in PAO1 and Iso 2.

| **Feature ID** | **Product** | **Fold change** | **P-value** | **Adjusted p-value n** | **PAO1 Means** | **Iso 2 Means** |
| --- | --- | --- | --- | --- | --- | --- |
| *algA* | phosphomannose isomerase / guanosine 5'-diphospho-D-mannose pyrophosphorylase | 20.08 | 2.11E-65 | 7.30E-64 | 39 | 1132 |
| *algD* | GDP-mannose 6-dehydrogenase AlgD | 15.91 | 3.78E-55 | 1.19E-53 | 45.5 | 1049.5 |
| *algF* | alginate o-acetyltransferase AlgF | 13.99 | 3.22E-47 | 9.57E-46 | 14.5 | 294.5 |
| *algX* | alginate biosynthesis protein AlgX | 13.82 | 2.54E-47 | 7.56E-46 | 13 | 260.5 |
| *algE* | Alginate production outer membrane protein AlgE precursor | 11.06 | 8.52E-52 | 2.65E-50 | 21.5 | 343 |
| *algJ* | alginate o-acetyltransferase AlgJ | 10.34 | 6.24E-26 | 1.29E-24 | 9.5 | 142.5 |
| *algL* | poly(beta-d-mannuronate) lyase precursor AlgL | 10.08 | 1.59E-46 | 4.68E-45 | 19 | 277 |
| *algK* | alginate biosynthetic protein AlgK precursor | 8.80 | 2.57E-26 | 5.39E-25 | 15.5 | 197.5 |
| *alg44* | alginate biosynthesis protein Alg44 | 8.27 | 7.19E-33 | 1.79E-31 | 22 | 263 |
| *alg8* | alginate biosynthesis protein Alg8 | 6.17 | 4.59E-39 | 1.23E-37 | 43 | 381.5 |
| *pelD* | PelD | 5.96 | 2.04E-39 | 5.54E-38 | 86.5 | 738 |
| *pelF* | PelF | 5.55 | 1.27E-39 | 3.47E-38 | 72 | 573 |
| *pelE* | PelE | 4.48 | 5.84E-20 | 9.53E-19 | 51.5 | 329.5 |
| PA4625 | cyclic diguanylate-regulated TPS partner A, CdrA | 4.37 | 4.24E-18 | 6.23E-17 | 956.5 | 6024.5 |
| *pelG* | PelG | 4.29 | 3.77E-19 | 5.93E-18 | 40.5 | 248.5 |
| *fliC* | flagellin type B | -7.77 | 0.00E+00 | 0.00E+00 | 41854.5 | 7728 |
| *flgL* | flagellar hook-associated protein type 3 FlgL | -8.03 | 2.01E-74 | 7.91E-73 | 1321 | 236.5 |
| *waaL* | O-antigen ligase, WaaL | -18.84 | 1.84E-18 | 2.74E-17 | 520.5 | 39.5 |
| *fgtA* | flagellar glycosyl transferase, FgtA | -67.33 | 5.84E-142 | 3.74E-140 | 4585.5 | 98 |
| *arr* | aminoglycoside response regulator, inner membrane phosphodiesterase | -249.99 | 1.32E-55 | 4.20E-54 | 202.5 | 1 |
| PA2771 | diguanylate cyclase with a self-blocked I-site, Dcsbis | -690.82 | 3.77E-16 | 4.87E-15 | 318.5 | 0.5 |
| *wbpL* | glycosyltransferase WbpL | -1048.06 | 1.72E-177 | 1.40E-175 | 1212.5 | 1.5 |
| *wbpH* | probable glycosyltransferase WbpH | -1239.27 | 9.90E-179 | 8.29E-177 | 2296.5 | 2.5 |
| *wbpJ* | probable glycosyl transferase WbpJ | -1334.51 | 1.05E-174 | 8.19E-173 | 1543.5 | 1.5 |
| *fliD* | flagellar capping protein FliD | -1940.68 | 8.78E-223 | 9.09E-221 | 6996 | 5 |
| *wbpE* | UDP-2-acetamido-2-dideoxy-d-ribo-hex-3-uluronic acid transaminase, wbpE | -2162.38 | 4.00E-185 | 3.73E-183 | 4751.5 | 3 |
| *wbpB* | UDP-2-acetamido-2-deoxy-d-glucuronic acid 3-dehydrogenase, WbpB | -2179.93 | 2.11E-176 | 1.70E-174 | 4038.5 | 2.5 |
| *wbpG* | LPS biosynthesis protein WbpG | -2295.22 | 1.54E-151 | 1.08E-149 | 4243 | 2.5 |
| *wbpA* | UDP-N-acetyl-d-glucosamine 6-Dehydrogenase | -2360.08 | 1.36E-216 | 1.39E-214 | 6851 | 4 |
| *wzz* | O-antigen chain length regulator | -2400.52 | 3.08E-166 | 2.37E-164 | 1103 | 0.5 |
| *wzx* | O-antigen translocase | -3664.63 | 1.74E-109 | 9.43E-108 | 406 | 0 |
| *wzy* | B-band O-antigen polymerase | -6615.39 | 3.66E-150 | 2.54E-148 | 732.5 | 0 |
| *wbpI* | UDP-N-acetylglucosamine 2-epimerase WbpI | -7555.15 | 1.56E-157 | 1.15E-155 | 3462 | 0.5 |
| *wbpK* | probable NAD-dependent epimerase/dehydratase WbpK | -10309.97 | 2.10E-162 | 1.60E-160 | 1140 | 0 |
| *wbpD* | UDP-2-acetamido-3-amino-2,3-dideoxy-d-glucuronic acid N-acetyltransferase, WbpD | -15416.88 | 8.61E-176 | 6.81E-174 | 1705.5 | 0 |
| *lasB* | elastase LasB | 1.62 | 1.46E-02 | 2.98E-02 | 4671 | 10885.5 |
| *rhlA* | rhamnosyltransferase chain A | 2.74 | 9.25E-16 | 1.16E-14 | 566.5 | 2227 |
| *rhlB* | rhamnosyltransferase chain B | 3.25 | 3.62E-19 | 5.71E-18 | 262 | 1222 |

## Conclusion

In this report, two SCVs of *P. aeruginosa* co-infecting with SARS-CoV-2 virus has been isolated from one critically illed COVID-19 patient and characterized at genomic, trancscriptomic and phenotypic levels. The isolates carry multiple specific GIs and several ARGs for survival, proliferation, drug-resistance and invasion in host’s respiratory systems. We demonstrated the alterations in biofilm-forming capability and QS expression in these SCVs during viral coinfection. Our results revealed the adaptations of *P. aeruginosa* to host respiratory system under the influence of SARS-CoV-2 during critical stage of COVID-19 infection. Understanding these adaptations may greatly contribute in the prognosis of disease development and treatment scheme decision to minimize the potential of secondary infections induced by *P. aeruginosa*.

## Note

### Author Contribution

JQ and ZC contributed to data analysis and manuscript preparation. YL(1), XKD, SH, YZ(1), ZJ, and YZ(2) contributed to experimental design and operation. CZ, YL(2), YL(3), LL and LY contributed to study design and manuscript verification.

### Funding

This work was supported by Guangdong Natural Science Foundation for Distinguished Young Scholar[2020B1515020003], Science and Technology Program of Guangzhou [201607020044], Science and Technology Program of Shenzhen [JCYJ20190809144005609] and Guangdong Basic and Applied Basic Research Foundation [2020A1515010586] for Dr. Jiuxin Qu, and Grant from Bill & Melinda Gates Foundation to Dr. Lei LIU.

### Ethics statement

This work is approved by the Ethics Committee of Shenzhen Third People’s Hospital, Second Hospital Affiliated to Southern University of science and Technology [2020-184].

