## Supplementary Materials and Methods for "Persistent bacterial coinfection of a COVID-19 patient caused by a genetically adapted *Pseudomonas aeruginosa* chronic colonizer"

**Antibiotic Susceptibility tests**

Two isolates of *Pseudomonas aeruginosa* were collected from sputum samples of one COVID-19 patient with 10 days interval (Isolate 1 on 12 February 2020 and Isolate 2 on 22 February 2020) during routine clinical tests. Bacterial antimicrobial susceptibility of the isolates with different colony morphologies, including ceftazidime, piperacillin, cefoperazone/sulbactam, imipenem, aztreonam, and levofloxacin, were performed using the Kirby-Bauer disc-diffusion method. Briefly, the isolates and reference strain were picked using sterile swab and adjusted to turbidity of 0.5 according to McFarland Standard in saline. A new sterile swab was used to absorb the bacterial suspension of each strain and spread plating evenly onto a MH agar plate(Detgerm, China). The plates were dried under room temperature. Antimicrobial susceptibility disks (ThermoFisher) were placed closely attaching to the surface of agar using sterile forceps. The plates were cultured for 16-18 hours at 35 ℃. Diameters of inhibition zone was meaured using a vernier caliper. Susceptibility was determined according to CLSI 2019.

**Bacterial strains and growth media**

The isolates and all used *P. aeruginosa* strains and plasmids are described in Table 1. Two *P. aeruginosa* isolates were collected from one Covid-19 patient and named Isolate 1(Iso1) and Isolate 2 (Iso 2), Cultures were grown in LB broth or ABTGC (0.1% MgCl_2_, 0.1% CaCl_2_, 0.1% FeCl_3,_ 10% A10 (15.1mM (NH4)_2_SO_4_, 33.7mM Na_2_HPO_4·_2H_2_O, 22.0 mM KH_2_PO_4_, 0.05mM NaCl), 0.2% glucose and 0.2% casamino acids) supplemented with 10% TSB. Antibiotic selection was performed with 300 μg/ml carbenicillin for *P. aeruginosa* strains carrying pUCP22-*cdrA::gfp* plasmid. Colony morphology of the isolates was imaged by Olympus(Mshot, China)with 8X magnification.

**Motility Assays**

Swimming motility of the isolates and *P. aeruginosa* PAO1 was tested on 0.3% LB agar plates. Swarming motility of the strains was tested on 0.5% agar plates supplemented with 0.8% nutrient broth and 0.5% glucose. Overnight cultures were diluted to OD_600nm_ of 0.01 and inoculated on agar plates using sterile toothpicks for swimming while 2 µl of the inoculums was spotted onto the surface of the agar plates for swarming. The plates were incubated at 37℃ statically for 14–16 h. Bacterial motility was imaged using Chemiluminescent(ChampChemiTM580,Sage,China)..

**Biofilm formation and crystal violet assay**

Overnight cultures of the isolates and PAO1 in LB medium were diluted to OD_600nm_ 0.01 in fresh LB broth. Triplicates of 100µL of the diluted cultures were aliquoted into 96-well microtiter plate. Cultures were then incubated statically for 24h at 37°C for biofilm formation. Spent medium was removed and wells were washed twice carefully with ddH_2_O. 125 µL of 0.1% crystal violet (CV) was added to each well and incubated for 15 min at room temperature. CV was removed and the wells were washed thoroughly twice with ddH_2_O. The plates were air-dried. CV stain was then dissolved into 125 µL of 30% acetic acid. Relative biofilm biomass was quantified by measuring optical density of CV staining on a Tecan infinity pro200 microplate reader at 550nm.

**QS inhibition Assay**

Overnight cultures of the isolates, PAO1, PAO1 quorum sensing (QS) mutant and reporter strains were diluted to OD_600nm_ 0.01 in ABTGC medium with 10%TSB and grown to OD_600nm_ 0.8~1.0. Then, cultures of the isolates, PAO1 and QS double mutant were centrifuged at 12,000g for 2 min to remove the cells and collect the supernatants. 50 µL of filtered cultural supernatants was collected and added to 50µL of *las* and *rhl* QS reporter strains respectively in triplicates. Green fluorescence and OD_600nm_ were recorded overnight (16h) in Tecan infinity pro200 microplate reader for QS expression of the strains. The expression of *las* and *rhl* QS was quantified by GFP/OD.

**Construction of cyclic-di-GMP reporter strains**

Plasmid pUCP22-*cdrA::gfp* was extracted from PAO1ΔPA0169/pUCP22-*cdrA::gfp*. The plasmid was transformed into PAO1 and the isolates by electroporation. The transformants were selected on LB agar plates containing 300µg/mL carbenicillin. The presence of the plasmid in the transformants was confirmed by extracting and running the plasmid on electrophoresis gel.

**Cyclic-di-GMP semi-quantification assay**

The transformants constructed above carrying pUCP22-*cdrA::gfp* reporter were cultivated in LB medium for overnight and diluted to OD_600nm_ 0.01 in ABTGC medium with 10%TSB. 100 μl of diluted bacteria were loaded into 96 well microplate in triplicates and allowed to grown to stationary phase. Green fluorescence and OD_600nm_ were measured using a Tecan Infinite Pro2000 microplate reader. Cyclic-di-GMP level expressed in the isolates and PAO1 was quantified by GFP/OD.

**Genome and transcriptome extraction**

Single colonies of the isolates were picked and grown in LB at 37°C, 200 rpm for 16 h. Genomic DNA of the isolates was extracted by AxyPerp Bacterial Genomic DNA Miniprep Kit (Corning, New York, USA) using manufacturer’s protocol. The PCR-free libraries of extracted genomic DNA were prepared by VAHTSTM PCR-Free DNA Library Prep Kit for Illumina®(Vazyme, China) following manufacturer’s protocol. Basically, DNA was fragmented and purifed by VAHTSTM DNA Clean Beads. Purified fragments were then tagged with VAHTSTM DNA Adapters for Illumina® (Vazyme, China). The concentrations and quality of the libraries were tested using qPCR and Agilent Technologies 2100 Bioanalyzer. The libraries were then sequenced on Illumina HiSeq X platform with paired end reads of 150 bp in length.

For transcriptomic sequencing, single colonies of the isolates were picked and grown in duplicates in LB at 37°C, 200 rpm until early stationary phase. Total RNA was extracted using Magen HiPure Universal RNA Mini kits (MCBio, China) according to the manufacturer’s instructions. RNA degradation and contamination were monitored on 1% agarose gels. The amount of RNA was measured by Qubit 2.0 (Thermo Fisher Scientific, MA, USA) and Nanodrop One (Thermo Fisher Scientific, MA, USA). RNA integrity was detected using Agilent 2100 system (Agilent Technologies, Waldbron, Germany). RNA libraries were prepared by NEB Next® Ultra™ Directional RNA Library Prep Kit for Illumina® (New England Biolabs, MA, USA) following manufacturer’s instruction. Ribosomal RNA were removed by Ribo-zero rRNA Removal Kit. Fragmentation was done using NEB Next First Strand Synthesis Reaction Buffer. cDNA was synthesized using random hexamer primer and M-MuLV Reverse Transcriptase (RNase H). cDNA fragments were selected with AMPure XP beads (Beckman Coulter, Beverly, USA). PCR products were then generated and purified with AMPure XP beads. The libraries were sequenced on an Illumina Novaseq platform with paired end reads of 150 bp in length.

For long read genomic sequencing, single colony of Iso 1 was picked and grown in LB at 37°C, 200 rpm for 16 h. Genomic DNA was extracted using Mabio Bacterial DNA Extraction Mini Kits (Mabio) according to the manufacturer’s instructions. DNA concentration, integrity and purity was monitored by the method described above. Qualified genomic DNA was fragmented with G-tubes (Covaris) and end-repaired to prepare SMRTbell DNA template libraries (with fragment size of >10 Kb selected by bluepippin system) according to the manufacturer’s specification (PacBio, Menlo Park, USA). Library quality was monitored by Qubit 3.0 Fluorometer (Life Technologies, Grand Island, NY) and average fragment size was estimated on Agilent 4200 system(Agilent, Santa Clara, CA). Genomic sequencing was then performed on the Pacific Biosciences RSII sequencer (PacBio, Menlo Park, USA) according to standard protocols.

**Sequencing data analysis**

Illumina genomic reads of the isolates was analyzed by CLC Genomics Workbench 20 (Qiagen) using Resequencing analysis module with 80% of frequency for single nucleotide polymorphism (SNP) of Iso 2 comparing to Iso 1 using PAO1 as reference genome.

RNA sequences of the two isolates and PAO1 were analyzed by RNA analysis module of CLC Genomics Workbench 20 (Qiagen) with default parameters. Significance of differential gene expression was determined based on absolute fold change≧4 and adjusted p-value<0.05. GO enrichment analysis of significantly expressed genes was performed on DAVID bioinformatics database v6.8 using EASE value of 0.05.[1] PCoA plot was drawn using vegan and ggplot2 packages in R software.

PacBio genomic reads of Iso 1 was assembled using HGAP4 pipeline using SMRT Link v9.0 with default settings. Multilocus sequence typing (MLST) of the assembled Iso1 genome was performed on the MLST server of Center of Genomic of Epidermiology using *P. aeruginosa* database.[2] Identification of antimicrobial resistance genes(ARGs) of Iso 1 was performed on the ResFinder 3.2 server of Center of Genomic of Epidermiology with minimal 85% of identity and minimal length of 60%.[3] Phylogenetic tree was drawn by libMUSCLE aligment mode of Parsnp package using whole genome sequences of the selected *P. aeruginosa* species. [4] Circular plot of selected genomes was perfomed using BLAST Ring Image Generator.[5] Genomic islands on Iso 1 were predicted by IslandViewer 4.[6] Genome annotation was done using RAST online annotation server using *P. aeruginosa* (Taxonomy ID 287) as reference.[7] All sequencing data are available under NCBI SRA BioProject XXX.
