## Supplementary Table S1 for "Persistent bacterial coinfection of a COVID-19 patient caused by a genetically adapted *Pseudomonas aeruginosa* chronic colonizer"

| **Strains or plasmids** | **Relevant genotype and/or characteristics** | **Reference** |
| --- | --- | --- |
| **strains** |  |  |
| *Pseudomonas aeruginosa* PAO1 | *Pseudomonas aeruginosa* ATCC | (1) |
| Isolate 1 (Iso 1) | Isolate from covid-19 patient | This study |
| Isolate 2 (Iso 2) | Isolate from covid-19 patient | This study |
| PAO1Δ*lasI*Δ*rhlI* | PAO1 *lasI* and *rhlI* double mutant | (2) |
| PAO1Δ*lasI*Δ*rhlI*/*P_lasB::gfp_* | PAO1 *lasI* and *rhlI* mutant containing *lasB-gfp*(ASV) reporter fusion |  |
| PAO1Δ*lasI*Δ*rhlI*/*P_rhlA::gfp_* | PAO1 *lasI* and *rhlI* mutant containing *rhlA-gfp*(ASV) reporter fusion | (3) |
| PAO1/*P_cdrA::gfp_* | PAO1 containing *cdrA-gfp* reporter fusion plasmid | This study |
| Iso 1/*P_cdrA::gfp_* | Isolate 1 containing *cdrA-gfp* reporter fusion plasmid | This study |
| Iso 2/*P_cdrA::gfp_* | Isolate 2 containing *cdrA-gfp* reporter fusion plasmid | This study |
| **plasmid** |  |  |
| pUCP22-*cdrA::gfp* |  | (4) |

**Table S1** **Bacteria strains and plasmids used in this study**

(1) Hentzer, M.; Riedel, K.; Rasmussen, T. B.; Heydorn, A.; Andersen, J. B.; Parsek, M. R.; Rice, S. A.; Eberl, L.; Molin, S.; Høiby, N.; Kjelleberg, S.; Givskov, M., Inhibition of quorum sensing in Pseudomonas aeruginosa biofilm bacteria by a halogenated furanone compound. Microbiology 2002, 148, 87-102.
(2) Hentzer, M.; Wu, H.; Andersen, J. B.; Riedel, K.; Rasmussen, T. B.; Bagge, N.; Kumar, N.; Schembri, M. A.; Song, Z.; Kristoffersen, P.; Manefield, M.; Costerton, J. W.; Molin, S.; Eberl, L.; Steinberg, P.; Kjelleberg, S.; Hoiby, N.; Givskov, M. Attenuation of Pseudomonas aeruginosa virulence by quorum sensing inhibitors. EMBO J. 2003, 22, 3803−3815.
(3) Hansen, S. K.; Rau, M. H.; Johansen, H. K.; Ciofu, O.; Jelsbak, L.; Yang, L.; Folkesson, A.; Jarmer, H. Ø.; Aanæs, K.; Buchwald, C. v.; Høiby, N.; Molin, S., Evolution and diversification of Pseudomonas aeruginosa in the paranasal sinuses of cystic fibrosis children have implications for chronic lung infection. ISME J. 2012, 6, 31-45.
(4) Rybtke MT, Borlee BR, Murakami K, Irie Y, Hentzer M, et al. 2012. Appl Environ Microbiol 78: 5060-9
