## Supplementary Table S2 for "Persistent bacterial coinfection of a COVID-19 patient caused by a genetically adapted *Pseudomonas aeruginosa* chronic colonizer"

| Date | Days of Hospitalization | WBC  (10^9/L) | Neutrophil  (10^9/L) | Lymphocyte  (10^9/L) | Interleukin-6  (pg/ml) | Antiviral Treatment | ANtimicrobial treatment | Antifungal treatment | Anti-inlfammatory treatment | Interferon | Immunity enhancement |
| --- | --- | --- | --- | --- | --- | --- | --- | --- | --- | --- | --- |
| 1/23/2020  (Admission) | Day 1 | NA | NA | NA | NA | Oseltamivir capsule (75mg) | Levofloxacin tablets(0.5g) |  |  | Recombinant human interferon α-1b injection (30ug) |  |
| 1/24/2020 | Day 2 | 9.14 | 7.34 | 1.42 | 28.16 | Oseltamivir capsule (75mg) | Levofloxacin tablets(0.5g) |  |  | Recombinant human interferon α-1b injection (30ug) |  |
| 1/25/2020 | Day 3 | 7.72 | 5.34 | 1.92 | NA | Oseltamivir capsule (75mg) | Levofloxacin tablets(0.5g) |  |  | Recombinant human interferon α-1b injection (30ug) |  |
| 1/26/2020 | Day 4 | 7.95 | 5.97 | 1.44 | 31.59 | lopinavir and ritonavir tablets (250mg) | Levofloxacin tablets(0.5g) |  | methylprednisolone injection(40mg) | Recombinant human interferon α-1b injection (60ug) | Immunoglobulin injection (2.5g) |
| 1/27/2020 | Day 5 | 11.51 | 8.74 | 1.88 | 30.07 | lopinavir and ritonavir tablets (251mg) | cefoperazone sodium and sulbactam sodium(1.5g) |  | methylprednisolone injection(40mg) | Recombinant human interferon α-1b injection (60ug) | Immunoglobulin injection (2.5g) |
| 1/28/2020 | Day 6 | 21.93 | 19.93 | 0.98 | 23.34 | lopinavir and ritonavir oral solution (160ml) | cefoperazone sodium and sulbactam sodium(3.0g),linezolid and glucose injection (600mg) |  | methylprednisolone injection(40mg) | Recombinant human interferon α-1b injection (60ug) | Immunoglobulin injection (2.5g),zadaxin (1.6mg) |
| 1/29/2020 | Day 7 | 30.23 | 27.70 | 1.17 | 8.38 | lopinavir and ritonavir oral solution (160ml) | Linezolid and glucose injection (600mg), cefoperazone sodium and sulbactam sodium injection (1.5g) |  |  | Recombinant human interferon α-1b injection (60ug) | Immunoglobulin injection (2.5g),zadaxin (1.6mg) |
| 1/30/2020 | Day 8 | 24.69 | 22.12 | 1.30 | 22.87 | lopinavir and ritonavir oral solution (160ml) | Linezolid and glucose injection (600mg), cefoperazone sodium and sulbactam sodium injection (1.5g) |  |  | Recombinant human interferon α-1b injection (60ug) | Immunoglobulin injection (2.5g),zadaxin (1.6mg) |
| 1/31/2020 | Day 9 | 14.91 | 12.63 | 1.865 | 62.38 | lopinavir and ritonavir oral solution (160ml) | Linezolid and glucose injection (600mg), cefoperazone sodium and sulbactam sodium injection (1.5g) |  |  | Recombinant human interferon α-1b injection (60ug) | Immunoglobulin injection (2.5g),zadaxin (1.6mg) |
| 2/1/2020 | Day 10 | 8.84 | 7.74 | 0.82 | NA | lopinavir and ritonavir oral solution (160ml) | Linezolid and glucose injection (600mg), cefoperazone sodium and sulbactam sodium injection (1.5g) |  |  | Recombinant human interferon α-1b injection (60ug) | Immunoglobulin injection (2.5g),zadaxin (1.6mg) |
| 2/2/2020 | Day 11 | 12.31 | 11.14 | 0.69 | NA | lopinavir and ritonavir oral solution (160ml) | Linezolid and glucose injection (600mg), cefoperazone sodium and sulbactam sodium injection (1.5g) |  |  | Recombinant human interferon α-1b injection (60ug) | Immunoglobulin injection (2.5g),zadaxin (1.6mg) |
| 2/3/2020 | Day 12 | 9.65 | 8.43 | 0.64 | NA | lopinavir and ritonavir oral solution (160ml) | Linezolid and glucose injection (600mg), cefoperazone sodium and sulbactam sodium injection (1.5g) |  |  | Recombinant human interferon α-1b injection (60ug) | Immunoglobulin injection (2.5g),zadaxin (1.6mg) |
| 2/4/2020 | Day 13 | 6.77 | 5.47 | 0.75 | NA | lopinavir and ritonavir oral solution (160ml) | Linezolid and glucose injection (600mg), cefoperazone sodium and sulbactam sodium injection (1.5g) |  |  | Recombinant human interferon α-1b injection (60ug) | Immunoglobulin injection (2.5g),zadaxin (1.6mg) |
| 2/5/2020 | Day 14 | 7.53 | 6.05 | 1.02 | NA | lopinavir and ritonavir oral solution (160ml) | Linezolid and glucose injection (600mg), cefoperazone sodium and sulbactam sodium injection (1.5g) | voriconazole tablets (200mg) |  | Recombinant human interferon α-1b injection (60ug) | Immunoglobulin injection (2.5g),zadaxin (1.6mg) |
| 2/6/2020 | Day 15 | 7.50 | 5.66 | 1.21 | NA | lopinavir and ritonavir oral solution (COVID-19) (1ml) | linezolid tablet (200mg),  cefoperazone sodium and sulbactam sodium(1.5g) | voriconazole tablets (200mg) |  | Recombinant human interferon α-1b injection (60ug) | Zadaxin (1.6mg) |
| 2/7/2020 | Day 16 | 8.39 | 6.45 | 1.39 | NA | lopinavir and ritonavir oral solution (COVID-19) (1ml) | cefoperazone sodium and sulbactam sodium(1.5g) | voriconazole tablets (200mg) |  | Recombinant human interferon α-1b injection (60ug) | Zadaxin (1.6mg) |
| 2/8/2020 | Day 17 | 6.85 | 4.69 | 1.53 | NA | lopinavir and ritonavir oral solution (COVID-19) (1ml) | cefoperazone sodium and sulbactam sodium(1.5g) | voriconazole injection (200mg) |  | Recombinant human interferon α-1b injection (60ug) | Zadaxin (1.6mg) |
| 2/9/2020 | Day 18 | 8.41 | 6.49 | 1.15 | NA | lopinavir and ritonavir oral solution (COVID-19) (1ml) | cefoperazone sodium and sulbactam sodium(1.5g) | voriconazole injection (200mg) |  | Recombinant human interferon α-1b injection (60ug) | Zadaxin (1.6mg) |
| 2/10/2020 | Day 19 | 6.55 | 4.72 | 1.34 | NA | lopinavir and ritonavir oral solution (COVID-19) (1ml) | cefoperazone sodium and sulbactam sodium(1.5g) | voriconazole injection (200mg) |  | Recombinant human interferon α-1b injection (60ug) | Zadaxin (1.6mg) |
| 2/11/2020 | Day 20 | 9.26 | 7.09 | 1.33 | NA | lopinavir and ritonavir oral solution (COVID-19) (1ml) | cefoperazone sodium and sulbactam sodium(1.5g) | voriconazole injection (200mg) |  | Recombinant human interferon α-1b injection (60ug) | Zadaxin (1.6mg) |
| 2/12/2020 | Day 21 | 12.73 | 9.89 | 1.97 | NA | lopinavir and ritonavir oral solution (COVID-19) (1ml) | Meropenem injection 0.5g | voriconazole injection (200mg) |  | Recombinant human interferon α-1b injection (60ug) | Zadaxin (1.6mg) |
| 2/13/2020 | Day 22 | 14.66 | 11.57 | 2.06 | NA | lopinavir and ritonavir oral solution (COVID-19) (1ml) | Meropenem injection 0.5g | voriconazole injection (200mg) |  | Recombinant human interferon α-1b injection (60ug) | Zadaxin (1.6mg) |
| 2/14/2020 | Day 23 | 13.54 | 10.10 | 2.03 | NA | lopinavir and ritonavir oral solution (COVID-19) (1ml) | Meropenem injection 0.5g | voriconazole injection (200mg) |  | Recombinant human interferon α-1b injection (60ug) |  |
| 2/15/2020 | Day 24 | 15.35 | 11.64 | 2.09 | 6.39 | lopinavir and ritonavir oral solution (COVID-19) (1ml) | Meropenem injection 0.5g | voriconazole injection (200mg) |  | Recombinant human interferon α-1b injection (60ug) |  |
| 2/16/2020 | Day 25 | 19.36 | 14.04 | 3.73 | 8.48 | Ganciclovir injection (0.25g) | Meropenem injection 0.5g |  |  | Recombinant human interferon α-1b injection (60ug) |  |
| 2/17/2020 | Day 26 | 14.92 | 10.43 | 2.97 | 5.95 | Ganciclovir injection (0.25g) | Meropenem injection 0.5g,Linezolid and glucose injection (600mg) |  |  | Recombinant human interferon α-1b injection (60ug) |  |
| 2/18/2020 | Day 27 | 13.54 | 9.56 | 2.82 | 5.27 | Ganciclovir injection (0.25g) | ceftazidime injection (1g) |  |  | Recombinant human interferon α-1b injection (60ug) |  |
| 2/19/2020 | Day 28 | 11.53 | 7.72 | 2.51 | 5.99 |  | ceftazidime injection (1g) |  |  | Recombinant human interferon α-1b injection (60ug) |  |
| 2/20/2020 | Day 29 | 13.12 | 9.06 | 2.74 | 6.28 |  | ceftazidime injection (1g) |  |  | Recombinant human interferon α-1b injection (60ug) |  |
| 2/21/2020 | Day 30 | 10.92 | 7.24 | 2.49 | 5.39 |  | ceftazidime injection (1g) |  |  | Recombinant human interferon α-1b injection (60ug) |  |
| 2/22/2020 | Day 31 | 10.29 | 7.13 | 2.10 | 5.10 |  | ceftazidime injection (1g), amikacin injection (0.2g) |  |  | Recombinant human interferon α-1b injection (60ug) |  |
| 2/23/2020 | Day 32 | 8.19 | 5.09 | 1.96 | 4.54 | lopinavir and ritonavir oral solution (COVID-19) (1ml) | ceftazidime injection (1g), amikacin injection (0.2g) |  |  | Recombinant human interferon α-1b injection (60ug) |  |
| 2/24/2020 | Day 33 | 8.39 | 5.45 | 1.94 | 4.16 | lopinavir and ritonavir oral solution (COVID-19) (1ml) | ceftazidime injection (1g), amikacin injection (0.2g) |  |  | Recombinant human interferon α-1b injection (60ug) |  |
| 2/25/2020 | Day 34 | NA | NA | NA | NA | lopinavir and ritonavir oral solution (COVID-19) (1ml) | ceftazidime injection (1g), amikacin injection (0.2g) |  |  | Recombinant human interferon α-1b injection (60ug) |  |
| 2/26/2020 | Day 35 | NA | NA | NA | NA | arbidol granules (0.1g) | ceftazidime injection (1g), amikacin injection (0.2g) |  |  | Recombinant human interferon α-1b injection (60ug) |  |
| 2/27/2020 | Day 36 | NA | NA | NA | NA | arbidol granules (0.1g) | ceftazidime injection (1g), amikacin injection (0.2g) |  |  |  |  |
| 2/28/2020 | Day 37 | NA | NA | NA | NA | arbidol granules (0.1g) | ceftazidime injection (1g), amikacin injection (0.2g) |  |  |  |  |
| 2/29/2020 | Day 38 | 6.56 | 3.52 | 2.27 | 2.59 | arbidol granules (0.1g) | ceftazidime injection (1g), amikacin injection (0.2g) |  |  |  |  |
| 3/1/2020 | Day 39 | NA | NA | NA | NA | arbidol granules (0.1g) | ceftazidime injection (1g), amikacin injection (0.2g) |  |  |  |  |
| 3/2/2020 | Day 40 | NA | NA | NA | NA | arbidol granules (0.1g) | ceftazidime injection (1g), amikacin injection (0.2g) |  |  |  |  |
| 3/3/2020 | Day 41 | 5.76 | 2.95 | 2.00 | <1.50 | arbidol granules (0.1g) | ceftazidime injection (1g), amikacin injection (0.2g) |  |  |  |  |
| 3/4/2020 | Day 42 | NA | NA | NA | NA | arbidol granules (0.1g) |  |  |  |  |  |
| 3/5/2020 | Day 43 | NA | NA | NA | NA | arbidol granules (0.1g) |  |  |  |  |  |
| 3/6/2020 | Day 44 | **4.83** | 2.15 | 2.07 | NA | arbidol granules (0.1g) |  |  |  |  |  |
| 3/7/2020  (Discharge) | Day 45 | NA | NA | NA | NA | arbidol granules (0.1g) |  |  |  |  |  |

**Table S2. Medical treatment schemes and blood index levels of WBC, N, L and IL-6 during hospitalization.**
