## Supplementary Table S3 for "Persistent bacterial coinfection of a COVID-19 patient caused by a genetically adapted *Pseudomonas aeruginosa* chronic colonizer"

| **Strain** | **GenBank Accession** |
| --- | --- |
| *Pseudomonas aeruginosa* DHS01 strain DH01 | NZ_CP013993.1 |
| *Pseudomonas aeruginosa* DK2 | NC_018080.1 |
| *Pseudomonas aeruginosa* strain F30658 | NZ_CP008857.1 |
| *Pseudomonas aeruginosa* strain H47921 | NZ_CP008861.1 |
| *Pseudomonas aeruginosa* strain LES431 | NC_023066.1 |
| *Pseudomonas aeruginosa* M18 | NC_017548.1 |
| *Pseudomonas aeruginosa* strain M1608 | NZ_CP008862.2 |
| *Pseudomonas aeruginosa* MTB-1 | NC_023019.1 |
| *Pseudomonas aeruginosa* strain N17-1 | NZ_CP014948.1 |
| *Pseudomonas aeruginosa* PA1 | NC_022808.2 |
| *Pseudomonas aeruginosa* RP73 | NC_021577.1 |
| *Pseudomonas aeruginosa* SCV20265 | NC_023149.1 |
| *Pseudomonas aeruginosa* SJTD-1 | NZ_CP015877.1 |
| *Pseudomonas aeruginosa* strain T63266 | NZ_CP008868.1 |
| *Pseudomonas aeruginosa* UCBPP-PA14 | NC_008463.1 |
| *Pseudomonas aeruginosa* YL84 | NZ_CP007147.1 |
| *Pseudomonas aeruginosa* strain Pa1207 | NZ_CP022001 |
| *Pseudomonas aeruginosa* strain PA8281 | NZ_CP015002 |
| *Pseudomonas aeruginosa* strain PAC1 | NZ_CP053706 |
| *Pseudomonas aeruginosa* strain PAO1611 | NZ_CP032126.1 |
| *Pseudomonas aeruginosa* strain PA_150577 | NZ_CP017306 |
| *Pseudomonas aeruginosa* strain PA34 | NZ_CP032552 |
| *Pseudomonas aeruginosa* PAO1 | NC_002516.2 |

Table S3. Genomic accession numbers of *P. aeruginosa* strains selected for phylogenetic tree construction
