## Supplementary Table S4 for "Persistent bacterial coinfection of a COVID-19 patient caused by a genetically adapted *Pseudomonas aeruginosa* chronic colonizer"

| **Island start** | **Island end** | **Length** | **Gene start** | **Gene end** | **Strand** | **Product** |
| --- | --- | --- | --- | --- | --- | --- |
| 135644 | 140361 | 4717 | 135644 | 136897 | 1 | MgtC family |
| 135644 | 140361 | 4717 | 136894 | 137154 | 1 | FIG00954700: hypothetical protein |
| 135644 | 140361 | 4717 | 137168 | 137563 | -1 | hypothetical protein |
| 135644 | 140361 | 4717 | 137762 | 137998 | 1 | hypothetical protein |
| 135644 | 140361 | 4717 | 138047 | 138517 | -1 | hypothetical protein |
| 135644 | 140361 | 4717 | 138933 | 139067 | -1 | hypothetical protein |
| 135644 | 140361 | 4717 | 139660 | 140361 | -1 | Transposase and inactivated derivatives |
| 199945 | 219302 | 19357 | 199945 | 201300 | 1 | hypothetical protein |
| 199945 | 219302 | 19357 | 201302 | 201619 | 1 | hypothetical protein |
| 199945 | 219302 | 19357 | 201623 | 202696 | 1 | hypothetical protein |
| 199945 | 219302 | 19357 | 203770 | 204633 | -1 | Integrase |
| 199945 | 219302 | 19357 | 205638 | 206810 | -1 | FIG141751: hypothetical protein in PFGI-1-like cluster |
| 199945 | 219302 | 19357 | 207698 | 207922 | 1 | hypothetical protein |
| 199945 | 219302 | 19357 | 208143 | 208430 | -1 | FIG00962252: hypothetical protein |
| 199945 | 219302 | 19357 | 208383 | 210275 | 1 | tRNA-5-carboxymethylaminomethyl-2- thiouridine(34) synthesis protein MnmG |
| 199945 | 219302 | 19357 | 210275 | 210919 | 1 | 16S rRNA (guanine(527)-N(7))-methyltransferase (EC 2.1.1.170) |
| 199945 | 219302 | 19357 | 210938 | 211726 | 1 | Chromosome (plasmid) partitioning protein ParA |
| 199945 | 219302 | 19357 | 211736 | 212608 | 1 | Chromosome (plasmid) partitioning protein ParB |
| 199945 | 219302 | 19357 | 212772 | 213152 | 1 | ATP synthase protein I |
| 199945 | 219302 | 19357 | 213169 | 214038 | 1 | ATP synthase F0 sector subunit a (EC 3.6.3.14) |
| 199945 | 219302 | 19357 | 214088 | 214345 | 1 | ATP synthase F0 sector subunit c (EC 3.6.3.14) |
| 199945 | 219302 | 19357 | 214403 | 214873 | 1 | ATP synthase F0 sector subunit b (EC 3.6.3.14) |
| 199945 | 219302 | 19357 | 214885 | 215421 | 1 | ATP synthase delta chain (EC 3.6.3.14) |
| 199945 | 219302 | 19357 | 215440 | 216984 | 1 | ATP synthase alpha chain (EC 3.6.3.14) |
| 199945 | 219302 | 19357 | 217035 | 217895 | 1 | ATP synthase gamma chain (EC 3.6.3.14) |
| 199945 | 219302 | 19357 | 217926 | 219302 | 1 | ATP synthase beta chain (EC 3.6.3.14) |
| 224566 | 240588 | 16022 | 224616 | 225443 | 1 | Transposon Tn7 transposition protein tnsA |
| 224566 | 240588 | 16022 | 225430 | 227523 | 1 | hypothetical protein |
| 224566 | 240588 | 16022 | 227516 | 229126 | 1 | Tn7-like transposition protein C |
| 224566 | 240588 | 16022 | 229128 | 230630 | 1 | Transposon Tn7 transposition protein tnsD |
| 224566 | 240588 | 16022 | 230623 | 232236 | 1 | hypothetical protein |
| 224566 | 240588 | 16022 | 232358 | 232549 | 1 | hypothetical protein |
| 224566 | 240588 | 16022 | 232619 | 235591 | 1 | Type I restriction-modification system, restriction subunit R (EC 3.1.21.3) |
| 224566 | 240588 | 16022 | 235588 | 235764 | 1 | ABC transporter |
| 224566 | 240588 | 16022 | 235761 | 237008 | 1 | Type I restriction-modification system, specificity subunit S |
| 224566 | 240588 | 16022 | 237008 | 238081 | 1 | Anticodon nuclease |
| 224566 | 240588 | 16022 | 238085 | 239719 | 1 | Type I restriction-modification system, DNA-methyltransferase subunit M (EC 2.1.1.72) |
| 224566 | 240588 | 16022 | 240135 | 240392 | 1 | hypothetical protein |
| 229128 | 239719 | 10591 | 229128 | 230630 | 1 | Transposon Tn7 transposition protein tnsD |
| 229128 | 239719 | 10591 | 230623 | 232236 | 1 | hypothetical protein |
| 229128 | 239719 | 10591 | 232358 | 232549 | 1 | hypothetical protein |
| 229128 | 239719 | 10591 | 232619 | 235591 | 1 | Type I restriction-modification system, restriction subunit R (EC 3.1.21.3) |
| 229128 | 239719 | 10591 | 235588 | 235764 | 1 | ABC transporter |
| 229128 | 239719 | 10591 | 235761 | 237008 | 1 | Type I restriction-modification system, specificity subunit S |
| 229128 | 239719 | 10591 | 237008 | 238081 | 1 | Anticodon nuclease |
| 229128 | 239719 | 10591 | 238085 | 239719 | 1 | Type I restriction-modification system, DNA-methyltransferase subunit M (EC 2.1.1.72) |
| 525128 | 545024 | 19896 | 525128 | 526177 | 1 | Phosphoserine phosphatase (EC 3.1.3.3) |
| 525128 | 545024 | 19896 | 526220 | 528205 | -1 | Secondary glycine betaine transporter BetU |
| 525128 | 545024 | 19896 | 528330 | 528794 | -1 | AAA+ ATPase superfamily protein YifB/ComM, associated with DNA recombination |
| 525128 | 545024 | 19896 | 528880 | 530976 | 1 | hypothetical protein |
| 525128 | 545024 | 19896 | 530976 | 532940 | 1 | hypothetical protein |
| 525128 | 545024 | 19896 | 533430 | 534788 | 1 | hypothetical protein |
| 525128 | 545024 | 19896 | 534942 | 536006 | -1 | ATP-dependent DNA helicase UvrD/PcrA (EC 3.6.4.12) |
| 525128 | 545024 | 19896 | 536181 | 536615 | -1 | Mercuric resistance operon regulatory protein MerR |
| 525128 | 545024 | 19896 | 536687 | 537037 | 1 | Mercuric transport protein, MerT |
| 525128 | 545024 | 19896 | 537015 | 537326 | 1 | Periplasmic mercury(+2) binding protein, MerP |
| 525128 | 545024 | 19896 | 537323 | 537784 | 1 | Mercuric transport protein, MerC |
| 525128 | 545024 | 19896 | 537836 | 539530 | 1 | Mercuric ion reductase (EC 1.16.1.1) |
| 525128 | 545024 | 19896 | 539548 | 539910 | 1 | Mercuric resistence transcriptional repressor, MerD |
| 525128 | 545024 | 19896 | 539907 | 540143 | 1 | Mercuric transport protein, MerE |
| 525128 | 545024 | 19896 | 540140 | 541018 | 1 | Tn21 protein of unknown function Urf2 |
| 525128 | 545024 | 19896 | 541103 | 542389 | -1 | Arsenite/antimonite:H+ antiporter ArsB |
| 525128 | 545024 | 19896 | 542875 | 544638 | -1 | Arsenite/antimonite pump-driving ATPase ArsA (EC 3.6.3.16) |
| 525128 | 545024 | 19896 | 544656 | 545024 | -1 | Asenic metallochaperone ArsD, transfers trivalent metalloids to ArsAB pump |
| 528795 | 541173 | 12378 | 528880 | 530976 | 1 | hypothetical protein |
| 528795 | 541173 | 12378 | 530976 | 532940 | 1 | hypothetical protein |
| 528795 | 541173 | 12378 | 533430 | 534788 | 1 | hypothetical protein |
| 528795 | 541173 | 12378 | 534942 | 536006 | -1 | ATP-dependent DNA helicase UvrD/PcrA (EC 3.6.4.12) |
| 528795 | 541173 | 12378 | 536181 | 536615 | -1 | Mercuric resistance operon regulatory protein MerR |
| 528795 | 541173 | 12378 | 536687 | 537037 | 1 | Mercuric transport protein, MerT |
| 528795 | 541173 | 12378 | 537015 | 537326 | 1 | Periplasmic mercury(+2) binding protein, MerP |
| 528795 | 541173 | 12378 | 537323 | 537784 | 1 | Mercuric transport protein, MerC |
| 528795 | 541173 | 12378 | 537836 | 539530 | 1 | Mercuric ion reductase (EC 1.16.1.1) |
| 528795 | 541173 | 12378 | 539548 | 539910 | 1 | Mercuric resistence transcriptional repressor, MerD |
| 528795 | 541173 | 12378 | 539907 | 540143 | 1 | Mercuric transport protein, MerE |
| 528795 | 541173 | 12378 | 540140 | 541018 | 1 | Tn21 protein of unknown function Urf2 |
| 528795 | 541173 | 12378 | 541103 | 542389 | -1 | Arsenite/antimonite:H+ antiporter ArsB |
| 528880 | 537326 | 8446 | 528880 | 530976 | 1 | hypothetical protein |
| 528880 | 537326 | 8446 | 530976 | 532940 | 1 | hypothetical protein |
| 528880 | 537326 | 8446 | 533430 | 534788 | 1 | hypothetical protein |
| 528880 | 537326 | 8446 | 534942 | 536006 | -1 | ATP-dependent DNA helicase UvrD/PcrA (EC 3.6.4.12) |
| 528880 | 537326 | 8446 | 536181 | 536615 | -1 | Mercuric resistance operon regulatory protein MerR |
| 528880 | 537326 | 8446 | 536687 | 537037 | 1 | Mercuric transport protein, MerT |
| 528880 | 537326 | 8446 | 537015 | 537326 | 1 | Periplasmic mercury(+2) binding protein, MerP |
| 528880 | 537326 | 8446 | 537323 | 537784 | 1 | Mercuric transport protein, MerC |
| 542201 | 546751 | 4550 | 541103 | 542389 | -1 | Arsenite/antimonite:H+ antiporter ArsB |
| 542201 | 546751 | 4550 | 542875 | 544638 | -1 | Arsenite/antimonite pump-driving ATPase ArsA (EC 3.6.3.16) |
| 542201 | 546751 | 4550 | 544656 | 545024 | -1 | Asenic metallochaperone ArsD, transfers trivalent metalloids to ArsAB pump |
| 542201 | 546751 | 4550 | 546103 | 546663 | 1 | Mobile element protein |
| 542201 | 546751 | 4550 | 546666 | 549632 | 1 | Mobile element protein |
| 549666 | 555613 | 5947 | 550174 | 552030 | -1 | ATP-dependent endonuclease family protein |
| 549666 | 555613 | 5947 | 552150 | 552965 | -1 | Sterol desaturase |
| 549666 | 555613 | 5947 | 552998 | 553759 | -1 | hypothetical protein |
| 549666 | 555613 | 5947 | 553818 | 554255 | -1 | hypothetical protein |
| 549666 | 555613 | 5947 | 554335 | 555567 | -1 | MFS-type efflux pump ArsJ specific for 1-arseno-3-phosphoglycerate |
| 549666 | 555613 | 5947 | 555585 | 556589 | -1 | NAD-dependent glyceraldehyde-3-phosphate dehydrogenase (EC 1.2.1.12) for arsenate detoxification |
| 550174 | 555567 | 5393 | 550174 | 552030 | -1 | ATP-dependent endonuclease family protein |
| 550174 | 555567 | 5393 | 552150 | 552965 | -1 | Sterol desaturase |
| 550174 | 555567 | 5393 | 552998 | 553759 | -1 | hypothetical protein |
| 550174 | 555567 | 5393 | 553818 | 554255 | -1 | hypothetical protein |
| 550174 | 555567 | 5393 | 554335 | 555567 | -1 | MFS-type efflux pump ArsJ specific for 1-arseno-3-phosphoglycerate |
| 559668 | 579439 | 19771 | 559428 | 559784 | -1 | Arsenical resistance operon repressor |
| 559668 | 579439 | 19771 | 559793 | 560206 | -1 | Arsenate reductase (EC 1.20.4.1) |
| 559668 | 579439 | 19771 | 560542 | 561339 | 1 | hypothetical protein |
| 559668 | 579439 | 19771 | 561460 | 563217 | -1 | Arsenite/antimonite pump-driving ATPase ArsA (EC 3.6.3.16) |
| 559668 | 579439 | 19771 | 563367 | 563498 | -1 | hypothetical protein |
| 559668 | 579439 | 19771 | 563720 | 565084 | -1 | putative secreted protein |
| 559668 | 579439 | 19771 | 565081 | 565596 | -1 | GCN5-related N-acetyltransferase |
| 559668 | 579439 | 19771 | 565658 | 566155 | -1 | transcriptional regulator, MarR family |
| 559668 | 579439 | 19771 | 566292 | 567488 | 1 | Uncharacterized MFS-type transporter |
| 559668 | 579439 | 19771 | 568018 | 568239 | -1 | hypothetical protein |
| 559668 | 579439 | 19771 | 568278 | 568490 | 1 | hypothetical protein |
| 559668 | 579439 | 19771 | 568774 | 568908 | 1 | hypothetical protein |
| 559668 | 579439 | 19771 | 568910 | 569554 | 1 | hypothetical protein |
| 559668 | 579439 | 19771 | 569949 | 570092 | -1 | hypothetical protein |
| 559668 | 579439 | 19771 | 570356 | 570511 | -1 | hypothetical protein |
| 559668 | 579439 | 19771 | 570596 | 571258 | -1 | hypothetical protein |
| 559668 | 579439 | 19771 | 571246 | 572205 | -1 | Glutathione synthase/Ribosomal protein S6 modification enzyme (glutaminyl transferase) |
| 559668 | 579439 | 19771 | 572234 | 572569 | -1 | hypothetical protein |
| 559668 | 579439 | 19771 | 572963 | 573277 | -1 | Mobile element protein |
| 559668 | 579439 | 19771 | 573891 | 574820 | 1 | Adenosine deaminase (EC 3.5.4.4) |
| 559668 | 579439 | 19771 | 575374 | 576168 | 1 | NAD-dependent protein deacetylases, SIR2 family |
| 559668 | 579439 | 19771 | 576354 | 576755 | 1 | hypothetical protein |
| 559668 | 579439 | 19771 | 576752 | 577036 | 1 | hypothetical protein |
| 559668 | 579439 | 19771 | 577148 | 577339 | 1 | hypothetical protein |
| 559668 | 579439 | 19771 | 577753 | 577983 | 1 | hypothetical protein |
| 559668 | 579439 | 19771 | 578046 | 578303 | 1 | hypothetical protein |
| 559668 | 579439 | 19771 | 578567 | 578788 | -1 | FIG00965560: hypothetical protein |
| 559668 | 579439 | 19771 | 579393 | 580466 | -1 | AAA+ ATPase superfamily protein YifB/ComM, associated with DNA recombination |
| 570356 | 578788 | 8432 | 570356 | 570511 | -1 | hypothetical protein |
| 570356 | 578788 | 8432 | 570596 | 571258 | -1 | hypothetical protein |
| 570356 | 578788 | 8432 | 571246 | 572205 | -1 | Glutathione synthase/Ribosomal protein S6 modification enzyme (glutaminyl transferase) |
| 570356 | 578788 | 8432 | 572234 | 572569 | -1 | hypothetical protein |
| 570356 | 578788 | 8432 | 572963 | 573277 | -1 | Mobile element protein |
| 570356 | 578788 | 8432 | 573891 | 574820 | 1 | Adenosine deaminase (EC 3.5.4.4) |
| 570356 | 578788 | 8432 | 575374 | 576168 | 1 | NAD-dependent protein deacetylases, SIR2 family |
| 570356 | 578788 | 8432 | 576354 | 576755 | 1 | hypothetical protein |
| 570356 | 578788 | 8432 | 576752 | 577036 | 1 | hypothetical protein |
| 570356 | 578788 | 8432 | 577148 | 577339 | 1 | hypothetical protein |
| 570356 | 578788 | 8432 | 577753 | 577983 | 1 | hypothetical protein |
| 570356 | 578788 | 8432 | 578046 | 578303 | 1 | hypothetical protein |
| 570356 | 578788 | 8432 | 578567 | 578788 | -1 | FIG00965560: hypothetical protein |
| 604315 | 608965 | 4650 | 603437 | 604318 | 1 | hypothetical protein |
| 604315 | 608965 | 4650 | 604315 | 607839 | 1 | FIG00954213: hypothetical protein |
| 604315 | 608965 | 4650 | 608105 | 608965 | 1 | hypothetical protein |
| 724484 | 733821 | 9337 | 724484 | 725365 | -1 | Glucose-1-phosphate thymidylyltransferase (EC 2.7.7.24) |
| 724484 | 733821 | 9337 | 725362 | 726270 | -1 | dTDP-4-dehydrorhamnose reductase (EC 1.1.1.133) |
| 724484 | 733821 | 9337 | 726267 | 727325 | -1 | dTDP-glucose 4,6-dehydratase (EC 4.2.1.46) |
| 724484 | 733821 | 9337 | 727401 | 727586 | 1 | hypothetical protein |
| 724484 | 733821 | 9337 | 727803 | 729662 | -1 | hypothetical protein |
| 724484 | 733821 | 9337 | 730120 | 731334 | -1 | hypothetical protein |
| 724484 | 733821 | 9337 | 732502 | 733821 | -1 | Integrase |
| 727401 | 733821 | 6420 | 727401 | 727586 | 1 | hypothetical protein |
| 727401 | 733821 | 6420 | 727803 | 729662 | -1 | hypothetical protein |
| 727401 | 733821 | 6420 | 730120 | 731334 | -1 | hypothetical protein |
| 727401 | 733821 | 6420 | 732502 | 733821 | -1 | Integrase |
| 840606 | 851241 | 10635 | 840606 | 840881 | 1 | hypothetical protein |
| 840606 | 851241 | 10635 | 840908 | 841735 | -1 | Transposase InsO for insertion sequence element IS911 |
| 840606 | 851241 | 10635 | 841762 | 842070 | -1 | Transposase InsN for insertion sequence element IS911 |
| 840606 | 851241 | 10635 | 842264 | 842680 | 1 | polyhydroxyalkanoate granule-associated protein PhaI |
| 840606 | 851241 | 10635 | 842691 | 843620 | 1 | Polyhydroxyalkanoate granule-associated protein PhaF |
| 840606 | 851241 | 10635 | 843664 | 844281 | -1 | Transcriptional regulator PhaD |
| 840606 | 851241 | 10635 | 844337 | 846019 | -1 | Polyhydroxyalkanoic acid synthase |
| 840606 | 851241 | 10635 | 846323 | 847180 | -1 | Poly(3-hydroxyalkanoate) depolymerase |
| 840606 | 851241 | 10635 | 847333 | 849012 | -1 | Polyhydroxyalkanoic acid synthase |
| 840606 | 851241 | 10635 | 849070 | 849285 | 1 | hypothetical protein |
| 840606 | 851241 | 10635 | 849432 | 849803 | -1 | FIG00899427: hypothetical protein |
| 840606 | 851241 | 10635 | 849898 | 851241 | -1 | ATP-dependent hsl protease ATP-binding subunit HslU |
| 1736627 | 1766258 | 29631 | 1735014 | 1736630 | 1 | probable chemotaxis transducer |
| 1736627 | 1766258 | 29631 | 1736627 | 1737832 | -1 | Chromate transport protein ChrA |
| 1736627 | 1766258 | 29631 | 1737887 | 1738690 | -1 | Transcriptional regulator, AraC family |
| 1736627 | 1766258 | 29631 | 1738790 | 1739680 | 1 | Permease of the drug/metabolite transporter (DMT) superfamily |
| 1736627 | 1766258 | 29631 | 1739793 | 1740476 | 1 | Lipoate-protein ligase A |
| 1736627 | 1766258 | 29631 | 1740517 | 1744032 | 1 | Exodeoxyribonuclease V gamma chain (EC 3.1.11.5) |
| 1736627 | 1766258 | 29631 | 1744029 | 1747766 | 1 | Exodeoxyribonuclease V beta chain (EC 3.1.11.5) |
| 1736627 | 1766258 | 29631 | 1747763 | 1749928 | 1 | Exodeoxyribonuclease V alpha chain (EC 3.1.11.5) |
| 1736627 | 1766258 | 29631 | 1749950 | 1753585 | -1 | Exonuclease SbcC |
| 1736627 | 1766258 | 29631 | 1753594 | 1754823 | -1 | Exonuclease SbcD |
| 1736627 | 1766258 | 29631 | 1761005 | 1764079 | -1 | Error-prone repair homolog of DNA polymerase III alpha subunit (EC 2.7.7.7) |
| 1736627 | 1766258 | 29631 | 1764076 | 1765491 | -1 | DNA polymerase IV-like protein ImuB |
| 1736627 | 1766258 | 29631 | 1765499 | 1766050 | -1 | RecA/RadA recombinase |
| 1736627 | 1766258 | 29631 | 1766142 | 1766258 | 1 | hypothetical protein |
| 1824319 | 1829976 | 5657 | 1824319 | 1824621 | 1 | Transcriptional regulator, Xre family |
| 1824319 | 1829976 | 5657 | 1824618 | 1825580 | 1 | hypothetical protein |
| 1824319 | 1829976 | 5657 | 1825596 | 1826489 | 1 | hypothetical protein |
| 1824319 | 1829976 | 5657 | 1826544 | 1828289 | 1 | hypothetical protein |
| 1824319 | 1829976 | 5657 | 1828349 | 1829200 | 1 | hypothetical protein |
| 1824319 | 1829976 | 5657 | 1829393 | 1829749 | 1 | Candidate type III effector Hop protein |
| 1824319 | 1829976 | 5657 | 1829746 | 1829976 | 1 | hypothetical protein |
| 1824504 | 1842149 | 17645 | 1824319 | 1824621 | 1 | Transcriptional regulator, Xre family |
| 1824504 | 1842149 | 17645 | 1824618 | 1825580 | 1 | hypothetical protein |
| 1824504 | 1842149 | 17645 | 1825596 | 1826489 | 1 | hypothetical protein |
| 1824504 | 1842149 | 17645 | 1826544 | 1828289 | 1 | hypothetical protein |
| 1824504 | 1842149 | 17645 | 1828349 | 1829200 | 1 | hypothetical protein |
| 1824504 | 1842149 | 17645 | 1829393 | 1829749 | 1 | Candidate type III effector Hop protein |
| 1824504 | 1842149 | 17645 | 1829746 | 1829976 | 1 | hypothetical protein |
| 1824504 | 1842149 | 17645 | 1830058 | 1830351 | 1 | hypothetical protein |
| 1824504 | 1842149 | 17645 | 1830361 | 1830756 | 1 | hypothetical protein |
| 1824504 | 1842149 | 17645 | 1830753 | 1831433 | 1 | hypothetical protein |
| 1824504 | 1842149 | 17645 | 1831430 | 1832317 | 1 | hypothetical protein |
| 1824504 | 1842149 | 17645 | 1832307 | 1833731 | 1 | hypothetical protein |
| 1824504 | 1842149 | 17645 | 1833712 | 1834146 | 1 | putative lipoprotein |
| 1824504 | 1842149 | 17645 | 1834259 | 1837027 | 1 | Type IV secretory pathway, VirB4 components |
| 1824504 | 1842149 | 17645 | 1837041 | 1837739 | 1 | Protein-disulfide isomerase |
| 1824504 | 1842149 | 17645 | 1838021 | 1838830 | 1 | putative protein Ymh |
| 1824504 | 1842149 | 17645 | 1838840 | 1839448 | -1 | hypothetical protein |
| 1824504 | 1842149 | 17645 | 1839511 | 1840449 | -1 | hypothetical protein |
| 1824504 | 1842149 | 17645 | 1840573 | 1841601 | -1 | hypothetical protein |
| 1830361 | 1837739 | 7378 | 1830361 | 1830756 | 1 | hypothetical protein |
| 1830361 | 1837739 | 7378 | 1830753 | 1831433 | 1 | hypothetical protein |
| 1830361 | 1837739 | 7378 | 1831430 | 1832317 | 1 | hypothetical protein |
| 1830361 | 1837739 | 7378 | 1832307 | 1833731 | 1 | hypothetical protein |
| 1830361 | 1837739 | 7378 | 1833712 | 1834146 | 1 | putative lipoprotein |
| 1830361 | 1837739 | 7378 | 1834259 | 1837027 | 1 | Type IV secretory pathway, VirB4 components |
| 1830361 | 1837739 | 7378 | 1837041 | 1837739 | 1 | Protein-disulfide isomerase |
| 1844465 | 1848479 | 4014 | 1843195 | 1844589 | 1 | hypothetical protein |
| 1844465 | 1848479 | 4014 | 1844586 | 1844915 | 1 | hypothetical protein |
| 1844465 | 1848479 | 4014 | 1845285 | 1845413 | 1 | hypothetical protein |
| 1844465 | 1848479 | 4014 | 1845460 | 1845849 | 1 | UPF0758 family protein |
| 1844465 | 1848479 | 4014 | 1846003 | 1846122 | -1 | hypothetical protein |
| 1844465 | 1848479 | 4014 | 1846433 | 1848280 | 1 | hypothetical protein |
| 1846433 | 1850974 | 4541 | 1846433 | 1848280 | 1 | hypothetical protein |
| 1846433 | 1850974 | 4541 | 1848497 | 1849783 | 1 | Serine/threonine protein kinase |
| 1846433 | 1850974 | 4541 | 1850009 | 1850974 | -1 | hypothetical protein |
| 1849719 | 1876816 | 27097 | 1848497 | 1849783 | 1 | Serine/threonine protein kinase |
| 1849719 | 1876816 | 27097 | 1850009 | 1850974 | -1 | hypothetical protein |
| 1849719 | 1876816 | 27097 | 1851501 | 1852700 | -1 | hypothetical protein |
| 1849719 | 1876816 | 27097 | 1852705 | 1853061 | -1 | hypothetical protein |
| 1849719 | 1876816 | 27097 | 1853063 | 1854418 | -1 | hypothetical protein |
| 1849719 | 1876816 | 27097 | 1854439 | 1856484 | -1 | VgrG protein |
| 1849719 | 1876816 | 27097 | 1856584 | 1857102 | -1 | T6SS component Hcp |
| 1849719 | 1876816 | 27097 | 1857468 | 1858343 | -1 | hypothetical protein |
| 1849719 | 1876816 | 27097 | 1858549 | 1858812 | -1 | hypothetical protein |
| 1849719 | 1876816 | 27097 | 1858809 | 1860005 | -1 | hypothetical protein |
| 1849719 | 1876816 | 27097 | 1860006 | 1862657 | -1 | hypothetical protein |
| 1849719 | 1876816 | 27097 | 1862685 | 1863554 | -1 | hypothetical protein |
| 1849719 | 1876816 | 27097 | 1863551 | 1865653 | -1 | VgrG protein |
| 1849719 | 1876816 | 27097 | 1865694 | 1866602 | -1 | hypothetical protein |
| 1849719 | 1876816 | 27097 | 1866604 | 1867941 | -1 | T6SS component TssK (ImpJ/VasE) |
| 1849719 | 1876816 | 27097 | 1867938 | 1868957 | -1 | Type VI secretion lipoprotein/VasD |
| 1849719 | 1876816 | 27097 | 1868957 | 1870321 | -1 | T6SS forkhead associated domain protein ImpI/VasC |
| 1849719 | 1876816 | 27097 | 1870318 | 1871337 | -1 | T6SS component TssG (ImpH/VasB) |
| 1849719 | 1876816 | 27097 | 1871301 | 1873067 | -1 | T6SS component TssF (ImpG/VasA) |
| 1849719 | 1876816 | 27097 | 1873089 | 1873517 | -1 | Uncharacterized protein similar to VCA0109 |
| 1849719 | 1876816 | 27097 | 1873526 | 1875007 | -1 | T6SS component TssC (ImpC/VipB) |
| 1849719 | 1876816 | 27097 | 1875195 | 1875698 | -1 | T6SS component TssB (ImpB/VipA) |
| 1849719 | 1876816 | 27097 | 1875894 | 1876028 | -1 | hypothetical protein |
| 1849719 | 1876816 | 27097 | 1876184 | 1878751 | 1 | T6SS AAA+ chaperone ClpV (TssH) |
| 1857468 | 1863554 | 6086 | 1857468 | 1858343 | -1 | hypothetical protein |
| 1857468 | 1863554 | 6086 | 1858549 | 1858812 | -1 | hypothetical protein |
| 1857468 | 1863554 | 6086 | 1858809 | 1860005 | -1 | hypothetical protein |
| 1857468 | 1863554 | 6086 | 1860006 | 1862657 | -1 | hypothetical protein |
| 1857468 | 1863554 | 6086 | 1862685 | 1863554 | -1 | hypothetical protein |
| 1857468 | 1863554 | 6086 | 1863551 | 1865653 | -1 | VgrG protein |
| 1879677 | 1892384 | 12707 | 1878766 | 1880340 | 1 | hypothetical protein |
| 1879677 | 1892384 | 12707 | 1880337 | 1881020 | 1 | Type VI secretion protein VasI |
| 1879677 | 1892384 | 12707 | 1881020 | 1882450 | 1 | hypothetical protein |
| 1879677 | 1892384 | 12707 | 1882466 | 1886203 | 1 | T6SS component TssM (IcmF/VasK) |
| 1879677 | 1892384 | 12707 | 1886229 | 1887266 | 1 | hypothetical protein |
| 1879677 | 1892384 | 12707 | 1887309 | 1889033 | 1 | hypothetical protein |
| 1879677 | 1892384 | 12707 | 1889791 | 1890186 | 1 | hypothetical protein |
| 1879677 | 1892384 | 12707 | 1890301 | 1891836 | -1 | Integrase |
| 1879677 | 1892384 | 12707 | 1891833 | 1892147 | -1 | hypothetical protein |
| 2156171 | 2161898 | 5727 | 2156171 | 2157451 | -1 | Integrase |
| 2156171 | 2161898 | 5727 | 2157448 | 2159367 | -1 | Pyruvate/2-oxoglutarate dehydrogenase complex, dihydrolipoamide acyltransferase (E2) component, and |
| 2156171 | 2161898 | 5727 | 2159702 | 2160835 | -1 | Putative oxidoreductase |
| 2156171 | 2161898 | 5727 | 2160978 | 2161898 | -1 | Transcriptional regulator, LysR family |
| 2163067 | 2167071 | 4004 | 2163067 | 2163702 | 1 | Putative oxidoreductase |
| 2163067 | 2167071 | 4004 | 2164060 | 2164410 | -1 | putative plasmid stablization protein |
| 2163067 | 2167071 | 4004 | 2164414 | 2164686 | -1 | Predicted transcriptional regulators containing the CopG/Arc/MetJ DNA-binding domain |
| 2163067 | 2167071 | 4004 | 2165201 | 2166727 | -1 | putative membrane protein |
| 2163067 | 2167071 | 4004 | 2166724 | 2167071 | -1 | FIG00953975: hypothetical protein |
| 2188295 | 2209868 | 21573 | 2188295 | 2188984 | -1 | hypothetical protein |
| 2188295 | 2209868 | 21573 | 2189306 | 2189557 | 1 | Transposase InsN for insertion sequence element IS911 |
| 2188295 | 2209868 | 21573 | 2189584 | 2190411 | 1 | Transposase InsO for insertion sequence element IS911 |
| 2188295 | 2209868 | 21573 | 2190520 | 2190660 | 1 | Mobile element protein |
| 2188295 | 2209868 | 21573 | 2191097 | 2192521 | -1 | hypothetical protein |
| 2188295 | 2209868 | 21573 | 2192521 | 2197563 | -1 | hypothetical protein |
| 2188295 | 2209868 | 21573 | 2197886 | 2198992 | 1 | Error-prone, lesion bypass DNA polymerase V (UmuC) |
| 2188295 | 2209868 | 21573 | 2199051 | 2200214 | 1 | hypothetical protein |
| 2188295 | 2209868 | 21573 | 2200245 | 2202494 | -1 | Superfamily II DNA/RNA helicases, SNF2 family |
| 2188295 | 2209868 | 21573 | 2202600 | 2204045 | -1 | FIG023873: Plasmid related protein |
| 2188295 | 2209868 | 21573 | 2204075 | 2204680 | -1 | FIG026997: Hypothetical protein |
| 2188295 | 2209868 | 21573 | 2204772 | 2205026 | -1 | FIG041301: Hypothetical protein |
| 2188295 | 2209868 | 21573 | 2205094 | 2205456 | -1 | FIG046709: Hypothetical protein |
| 2188295 | 2209868 | 21573 | 2205559 | 2206350 | -1 | FIG034376: Hypothetical protein |
| 2188295 | 2209868 | 21573 | 2206407 | 2206757 | -1 | FIG00960315: hypothetical protein |
| 2188295 | 2209868 | 21573 | 2206993 | 2207700 | -1 | FIG00902157: hypothetical protein |
| 2188295 | 2209868 | 21573 | 2207898 | 2208050 | 1 | hypothetical protein |
| 2188295 | 2209868 | 21573 | 2208185 | 2208382 | -1 | hypothetical protein |
| 2188295 | 2209868 | 21573 | 2208456 | 2208614 | -1 | FIG00955915: hypothetical protein |
| 2188295 | 2209868 | 21573 | 2208752 | 2209231 | -1 | FIG051360: Periplasmic protein TonB, links inner and outer membranes |
| 2188295 | 2209868 | 21573 | 2209556 | 2209690 | -1 | hypothetical protein |
| 2188295 | 2209868 | 21573 | 2209692 | 2209868 | -1 | hypothetical protein |
| 2189584 | 2197563 | 7979 | 2189584 | 2190411 | 1 | Transposase InsO for insertion sequence element IS911 |
| 2189584 | 2197563 | 7979 | 2190520 | 2190660 | 1 | Mobile element protein |
| 2189584 | 2197563 | 7979 | 2191097 | 2192521 | -1 | hypothetical protein |
| 2189584 | 2197563 | 7979 | 2192521 | 2197563 | -1 | hypothetical protein |
| 2201522 | 2237706 | 36184 | 2200245 | 2202494 | -1 | Superfamily II DNA/RNA helicases, SNF2 family |
| 2201522 | 2237706 | 36184 | 2202600 | 2204045 | -1 | FIG023873: Plasmid related protein |
| 2201522 | 2237706 | 36184 | 2204075 | 2204680 | -1 | FIG026997: Hypothetical protein |
| 2201522 | 2237706 | 36184 | 2204772 | 2205026 | -1 | FIG041301: Hypothetical protein |
| 2201522 | 2237706 | 36184 | 2205094 | 2205456 | -1 | FIG046709: Hypothetical protein |
| 2201522 | 2237706 | 36184 | 2205559 | 2206350 | -1 | FIG034376: Hypothetical protein |
| 2201522 | 2237706 | 36184 | 2206407 | 2206757 | -1 | FIG00960315: hypothetical protein |
| 2201522 | 2237706 | 36184 | 2206993 | 2207700 | -1 | FIG00902157: hypothetical protein |
| 2201522 | 2237706 | 36184 | 2207898 | 2208050 | 1 | hypothetical protein |
| 2201522 | 2237706 | 36184 | 2208185 | 2208382 | -1 | hypothetical protein |
| 2201522 | 2237706 | 36184 | 2208456 | 2208614 | -1 | FIG00955915: hypothetical protein |
| 2201522 | 2237706 | 36184 | 2208752 | 2209231 | -1 | FIG051360: Periplasmic protein TonB, links inner and outer membranes |
| 2201522 | 2237706 | 36184 | 2209556 | 2209690 | -1 | hypothetical protein |
| 2201522 | 2237706 | 36184 | 2209692 | 2209868 | -1 | hypothetical protein |
| 2201522 | 2237706 | 36184 | 2209943 | 2210188 | -1 | FIG00963899: hypothetical protein |
| 2201522 | 2237706 | 36184 | 2211254 | 2211691 | -1 | Conjugative transfer protein PilM in PFGI-1-like cluster |
| 2201522 | 2237706 | 36184 | 2211720 | 2213048 | -1 | hypothetical protein |
| 2201522 | 2237706 | 36184 | 2213053 | 2213994 | -1 | Conjugative transfer ATPase PilU in PFGI-1-like cluster |
| 2201522 | 2237706 | 36184 | 2213991 | 2214521 | -1 | Conjugative transfer protein PilS in PFGI-1-like cluster |
| 2201522 | 2237706 | 36184 | 2214543 | 2215622 | -1 | hypothetical protein |
| 2201522 | 2237706 | 36184 | 2215622 | 2217202 | -1 | IncI1 plasmid conjugative transfer ATPase PilQ |
| 2201522 | 2237706 | 36184 | 2217211 | 2217744 | -1 | Conjugative transfer protein PilP in PFGI-1-like cluster |
| 2201522 | 2237706 | 36184 | 2217734 | 2219059 | -1 | hypothetical protein |
| 2201522 | 2237706 | 36184 | 2219063 | 2220772 | -1 | Conjugative transfer protein PilN in PFGI-1-like cluster |
| 2201522 | 2237706 | 36184 | 2220772 | 2221896 | -1 | hypothetical protein |
| 2201522 | 2237706 | 36184 | 2222281 | 2222574 | 1 | hypothetical protein |
| 2201522 | 2237706 | 36184 | 2222655 | 2223566 | -1 | Hydrolase, alpha/beta fold family |
| 2201522 | 2237706 | 36184 | 2223616 | 2224542 | -1 | hypothetical protein |
| 2201522 | 2237706 | 36184 | 2224641 | 2225414 | -1 | 2,3-dihydroxy-2,3-dihydro-phenylpropionate dehydrogenase (EC 1.3.1.-) |
| 2201522 | 2237706 | 36184 | 2225440 | 2226906 | -1 | Aldehyde dehydrogenase (EC 1.2.1.3) |
| 2201522 | 2237706 | 36184 | 2227321 | 2227509 | -1 | hypothetical protein |
| 2201522 | 2237706 | 36184 | 2227595 | 2229568 | -1 | hypothetical protein |
| 2201522 | 2237706 | 36184 | 2229565 | 2231454 | -1 | hypothetical protein |
| 2201522 | 2237706 | 36184 | 2231618 | 2232523 | -1 | hypothetical protein |
| 2201522 | 2237706 | 36184 | 2232641 | 2233393 | -1 | Haemophilus-specific protein, uncharacterized |
| 2201522 | 2237706 | 36184 | 2233402 | 2235321 | -1 | DNA topoisomerase I (EC 5.99.1.2) |
| 2201522 | 2237706 | 36184 | 2235677 | 2236165 | -1 | Single-stranded DNA-binding protein in PFGI-1-like cluster |
| 2201522 | 2237706 | 36184 | 2236179 | 2236712 | -1 | Integrase regulator R |
| 2201522 | 2237706 | 36184 | 2236718 | 2237446 | -1 | FIG141694: hypothetical protein in PFGI-1-like cluster |
| 2201522 | 2237706 | 36184 | 2237603 | 2237719 | -1 | FIG00960543: hypothetical protein |
| 2222281 | 2227509 | 5228 | 2222281 | 2222574 | 1 | hypothetical protein |
| 2222281 | 2227509 | 5228 | 2222655 | 2223566 | -1 | Hydrolase, alpha/beta fold family |
| 2222281 | 2227509 | 5228 | 2223616 | 2224542 | -1 | hypothetical protein |
| 2222281 | 2227509 | 5228 | 2224641 | 2225414 | -1 | 2,3-dihydroxy-2,3-dihydro-phenylpropionate dehydrogenase (EC 1.3.1.-) |
| 2222281 | 2227509 | 5228 | 2225440 | 2226906 | -1 | Aldehyde dehydrogenase (EC 1.2.1.3) |
| 2222281 | 2227509 | 5228 | 2227321 | 2227509 | -1 | hypothetical protein |
| 2238878 | 2243837 | 4959 | 2238878 | 2239039 | -1 | FIG00960543: hypothetical protein |
| 2238878 | 2243837 | 4959 | 2239437 | 2239751 | -1 | FIG00960543: hypothetical protein |
| 2238878 | 2243837 | 4959 | 2239748 | 2241073 | -1 | FIG141751: hypothetical protein in PFGI-1-like cluster |
| 2238878 | 2243837 | 4959 | 2241070 | 2241825 | -1 | FIG004780: hypothetical protein in PFGI-1-like cluster |
| 2238878 | 2243837 | 4959 | 2241853 | 2243580 | -1 | Protein with ParB-like nuclease domain in PFGI-1-like cluster |
| 2238878 | 2243837 | 4959 | 2243583 | 2243837 | -1 | FIG00960798: hypothetical protein |
| 2238945 | 2243915 | 4970 | 2238878 | 2239039 | -1 | FIG00960543: hypothetical protein |
| 2238945 | 2243915 | 4970 | 2239437 | 2239751 | -1 | FIG00960543: hypothetical protein |
| 2238945 | 2243915 | 4970 | 2239748 | 2241073 | -1 | FIG141751: hypothetical protein in PFGI-1-like cluster |
| 2238945 | 2243915 | 4970 | 2241070 | 2241825 | -1 | FIG004780: hypothetical protein in PFGI-1-like cluster |
| 2238945 | 2243915 | 4970 | 2241853 | 2243580 | -1 | Protein with ParB-like nuclease domain in PFGI-1-like cluster |
| 2238945 | 2243915 | 4970 | 2243583 | 2243837 | -1 | FIG00960798: hypothetical protein |
| 2238945 | 2243915 | 4970 | 2243840 | 2244856 | -1 | Nucleoid-associated protein NdpA |
| 2241070 | 2263262 | 22192 | 2239748 | 2241073 | -1 | FIG141751: hypothetical protein in PFGI-1-like cluster |
| 2241070 | 2263262 | 22192 | 2241070 | 2241825 | -1 | FIG004780: hypothetical protein in PFGI-1-like cluster |
| 2241070 | 2263262 | 22192 | 2241853 | 2243580 | -1 | Protein with ParB-like nuclease domain in PFGI-1-like cluster |
| 2241070 | 2263262 | 22192 | 2243583 | 2243837 | -1 | FIG00960798: hypothetical protein |
| 2241070 | 2263262 | 22192 | 2243840 | 2244856 | -1 | Nucleoid-associated protein NdpA |
| 2241070 | 2263262 | 22192 | 2244856 | 2245089 | -1 | FIG034647: hypothetical protein in PFGI-1-like cluster |
| 2241070 | 2263262 | 22192 | 2245082 | 2245255 | -1 | FIG00954464: hypothetical protein |
| 2241070 | 2263262 | 22192 | 2245569 | 2245826 | -1 | FIG00963725: hypothetical protein |
| 2241070 | 2263262 | 22192 | 2245823 | 2246350 | -1 | FIG00957722: hypothetical protein |
| 2241070 | 2263262 | 22192 | 2246340 | 2246525 | -1 | hypothetical protein |
| 2241070 | 2263262 | 22192 | 2246932 | 2247891 | -1 | hypothetical protein |
| 2241070 | 2263262 | 22192 | 2247948 | 2249291 | -1 | Replicative DNA helicase (DnaB) (EC 3.6.4.12) |
| 2241070 | 2263262 | 22192 | 2249288 | 2249506 | -1 | FIG00953473: hypothetical protein |
| 2241070 | 2263262 | 22192 | 2249490 | 2250197 | -1 | Phage protein |
| 2241070 | 2263262 | 22192 | 2250194 | 2250895 | -1 | FIG004780: hypothetical protein in PFGI-1-like cluster |
| 2241070 | 2263262 | 22192 | 2250895 | 2251341 | -1 | FIG00955294: hypothetical protein |
| 2241070 | 2263262 | 22192 | 2251578 | 2252333 | -1 | FIG00954566: hypothetical protein |
| 2241070 | 2263262 | 22192 | 2252330 | 2252827 | -1 | hypothetical protein |
| 2241070 | 2263262 | 22192 | 2252824 | 2253561 | -1 | Orf50 |
| 2241070 | 2263262 | 22192 | 2253563 | 2254429 | -1 | ParA-like protein |
| 2241070 | 2263262 | 22192 | 2254934 | 2256217 | -1 | Integrase |
| 2241070 | 2263262 | 22192 | 2256214 | 2258133 | -1 | Pyruvate/2-oxoglutarate dehydrogenase complex, dihydrolipoamide acyltransferase (E2) component, and |
| 2241070 | 2263262 | 22192 | 2258824 | 2259693 | 1 | bacteriocin, putative |
| 2241070 | 2263262 | 22192 | 2260238 | 2260588 | -1 | putative plasmid stablization protein |
| 2241070 | 2263262 | 22192 | 2260592 | 2260864 | -1 | Predicted transcriptional regulators containing the CopG/Arc/MetJ DNA-binding domain |
| 2241070 | 2263262 | 22192 | 2260983 | 2261333 | 1 | hypothetical protein |
| 2241070 | 2263262 | 22192 | 2261368 | 2262912 | -1 | putative membrane protein |
| 2241070 | 2263262 | 22192 | 2262909 | 2263262 | -1 | FIG00953975: hypothetical protein |
| 2244083 | 2254690 | 10607 | 2243840 | 2244856 | -1 | Nucleoid-associated protein NdpA |
| 2244083 | 2254690 | 10607 | 2244856 | 2245089 | -1 | FIG034647: hypothetical protein in PFGI-1-like cluster |
| 2244083 | 2254690 | 10607 | 2245082 | 2245255 | -1 | FIG00954464: hypothetical protein |
| 2244083 | 2254690 | 10607 | 2245569 | 2245826 | -1 | FIG00963725: hypothetical protein |
| 2244083 | 2254690 | 10607 | 2245823 | 2246350 | -1 | FIG00957722: hypothetical protein |
| 2244083 | 2254690 | 10607 | 2246340 | 2246525 | -1 | hypothetical protein |
| 2244083 | 2254690 | 10607 | 2246932 | 2247891 | -1 | hypothetical protein |
| 2244083 | 2254690 | 10607 | 2247948 | 2249291 | -1 | Replicative DNA helicase (DnaB) (EC 3.6.4.12) |
| 2244083 | 2254690 | 10607 | 2249288 | 2249506 | -1 | FIG00953473: hypothetical protein |
| 2244083 | 2254690 | 10607 | 2249490 | 2250197 | -1 | Phage protein |
| 2244083 | 2254690 | 10607 | 2250194 | 2250895 | -1 | FIG004780: hypothetical protein in PFGI-1-like cluster |
| 2244083 | 2254690 | 10607 | 2250895 | 2251341 | -1 | FIG00955294: hypothetical protein |
| 2244083 | 2254690 | 10607 | 2251578 | 2252333 | -1 | FIG00954566: hypothetical protein |
| 2244083 | 2254690 | 10607 | 2252330 | 2252827 | -1 | hypothetical protein |
| 2244083 | 2254690 | 10607 | 2252824 | 2253561 | -1 | Orf50 |
| 2244083 | 2254690 | 10607 | 2253563 | 2254429 | -1 | ParA-like protein |
| 2252824 | 2258133 | 5309 | 2252330 | 2252827 | -1 | hypothetical protein |
| 2252824 | 2258133 | 5309 | 2252824 | 2253561 | -1 | Orf50 |
| 2252824 | 2258133 | 5309 | 2253563 | 2254429 | -1 | ParA-like protein |
| 2252824 | 2258133 | 5309 | 2254934 | 2256217 | -1 | Integrase |
| 2252824 | 2258133 | 5309 | 2256214 | 2258133 | -1 | Pyruvate/2-oxoglutarate dehydrogenase complex, dihydrolipoamide acyltransferase (E2) component, and |
| 2272358 | 2277019 | 4661 | 2272530 | 2273114 | -1 | RNA polymerase ECF-type sigma factor |
| 2272358 | 2277019 | 4661 | 2274543 | 2274881 | 1 | FIG00956135: hypothetical protein |
| 2272358 | 2277019 | 4661 | 2275299 | 2276723 | 1 | Coproporphyrinogen III oxidase, oxygen-independent (EC 1.3.99.22) |
| 2272358 | 2277019 | 4661 | 2276725 | 2278293 | -1 | Anaerobic nitric oxide reductase transcription regulator NorR |
| 2274543 | 2296162 | 21619 | 2274543 | 2274881 | 1 | FIG00956135: hypothetical protein |
| 2274543 | 2296162 | 21619 | 2275299 | 2276723 | 1 | Coproporphyrinogen III oxidase, oxygen-independent (EC 1.3.99.22) |
| 2274543 | 2296162 | 21619 | 2276725 | 2278293 | -1 | Anaerobic nitric oxide reductase transcription regulator NorR |
| 2274543 | 2296162 | 21619 | 2278392 | 2278493 | 1 | hypothetical protein |
| 2274543 | 2296162 | 21619 | 2278486 | 2280762 | 1 | Nitric-oxide reductase (EC 1.7.99.7), quinol-dependent |
| 2274543 | 2296162 | 21619 | 2281226 | 2281366 | 1 | Transposase InsO for insertion sequence element IS911 |
| 2274543 | 2296162 | 21619 | 2281543 | 2282208 | -1 | hypothetical protein |
| 2274543 | 2296162 | 21619 | 2282315 | 2282788 | 1 | Mercuric resistance operon regulatory protein MerR |
| 2274543 | 2296162 | 21619 | 2284253 | 2285359 | 1 | NADH:flavin oxidoreductases, Old Yellow Enzyme family |
| 2274543 | 2296162 | 21619 | 2285511 | 2285627 | 1 | hypothetical protein |
| 2274543 | 2296162 | 21619 | 2285921 | 2286325 | 1 | hypothetical protein |
| 2274543 | 2296162 | 21619 | 2286446 | 2287006 | -1 | hypothetical protein |
| 2274543 | 2296162 | 21619 | 2287384 | 2287695 | -1 | Phosphohydrolase (MutT/nudix family protein) |
| 2274543 | 2296162 | 21619 | 2287677 | 2288087 | -1 | Phosphohydrolase (MutT/nudix family protein) |
| 2274543 | 2296162 | 21619 | 2288283 | 2288924 | -1 | hypothetical protein |
| 2274543 | 2296162 | 21619 | 2289442 | 2290059 | 1 | Fimbrial protein precursor |
| 2274543 | 2296162 | 21619 | 2290117 | 2290830 | 1 | hypothetical protein |
| 2274543 | 2296162 | 21619 | 2290931 | 2293450 | 1 | hypothetical protein |
| 2274543 | 2296162 | 21619 | 2293530 | 2294051 | -1 | Methylated-DNA--protein-cysteine methyltransferase (EC 2.1.1.63) |
| 2274543 | 2296162 | 21619 | 2294609 | 2296162 | 1 | PQS biosynthesis protein PqsA, anthranilate-CoA ligase (EC 6.2.1.32) |
| 2274543 | 2296162 | 21619 | 2296156 | 2297007 | 1 | PQS biosynthesis protein PqsB, similar to 3-oxoacyl-[acyl-carrier-protein] synthase III |
| 2277620 | 2282783 | 5163 | 2276725 | 2278293 | -1 | Anaerobic nitric oxide reductase transcription regulator NorR |
| 2277620 | 2282783 | 5163 | 2278392 | 2278493 | 1 | hypothetical protein |
| 2277620 | 2282783 | 5163 | 2278486 | 2280762 | 1 | Nitric-oxide reductase (EC 1.7.99.7), quinol-dependent |
| 2277620 | 2282783 | 5163 | 2281226 | 2281366 | 1 | Transposase InsO for insertion sequence element IS911 |
| 2277620 | 2282783 | 5163 | 2281543 | 2282208 | -1 | hypothetical protein |
| 2277620 | 2282783 | 5163 | 2282315 | 2282788 | 1 | Mercuric resistance operon regulatory protein MerR |
| 2464593 | 2479194 | 14601 | 2464593 | 2464961 | -1 | hypothetical protein |
| 2464593 | 2479194 | 14601 | 2465389 | 2473647 | -1 | Alkaline phosphatase (EC 3.1.3.1) |
| 2464593 | 2479194 | 14601 | 2474355 | 2475770 | -1 | RTX toxin transporter, determinant D |
| 2464593 | 2479194 | 14601 | 2475767 | 2477923 | -1 | RTX toxin transporter, ATP-binding protein |
| 2464593 | 2479194 | 14601 | 2477968 | 2479194 | -1 | Putative glycosyltransferase |
| 2714176 | 2727726 | 13550 | 2714176 | 2715189 | 1 | UDP-glucose 4-epimerase (EC 5.1.3.2) |
| 2714176 | 2727726 | 13550 | 2715186 | 2716337 | 1 | Putative glycosyl transferase |
| 2714176 | 2727726 | 13550 | 2716639 | 2717625 | 1 | Export ABC transporter, ATP-binding protein |
| 2714176 | 2727726 | 13550 | 2718102 | 2719328 | 1 | hypothetical protein |
| 2714176 | 2727726 | 13550 | 2719345 | 2720073 | 1 | hypothetical protein |
| 2714176 | 2727726 | 13550 | 2720170 | 2721654 | 1 | Putative glycosyl transferase |
| 2714176 | 2727726 | 13550 | 2721651 | 2722874 | 1 | Putative glycosyl transferase |
| 2714176 | 2727726 | 13550 | 2722871 | 2724643 | 1 | Putative glycosyl transferase |
| 2714176 | 2727726 | 13550 | 2724665 | 2725888 | 1 | hypothetical protein |
| 2714176 | 2727726 | 13550 | 2725945 | 2726535 | -1 | Adenylylsulfate kinase (EC 2.7.1.25) |
| 2714176 | 2727726 | 13550 | 2727040 | 2727294 | 1 | hypothetical protein |
| 2714176 | 2727726 | 13550 | 2727310 | 2727726 | 1 | hypothetical protein |
| 2861569 | 2871301 | 9732 | 2861569 | 2862000 | 1 | GlcG protein |
| 2861569 | 2871301 | 9732 | 2862404 | 2863054 | -1 | Transcriptional regulator, AcrR family |
| 2861569 | 2871301 | 9732 | 2863290 | 2864285 | 1 | GTP 3',8-cyclase (EC 4.1.99.22) |
| 2861569 | 2871301 | 9732 | 2864470 | 2864799 | 1 | Putative phosphatase |
| 2861569 | 2871301 | 9732 | 2864891 | 2866168 | -1 | Xanthine/uracil permease family protein |
| 2861569 | 2871301 | 9732 | 2866169 | 2866447 | 1 | FIG00954510: hypothetical protein |
| 2861569 | 2871301 | 9732 | 2866941 | 2867201 | -1 | T6SS PAAR-repeat protein |
| 2861569 | 2871301 | 9732 | 2867314 | 2867622 | 1 | Transposase InsN for insertion sequence element IS911 |
| 2861569 | 2871301 | 9732 | 2867649 | 2868476 | 1 | Transposase InsO for insertion sequence element IS911 |
| 2861569 | 2871301 | 9732 | 2868480 | 2869595 | -1 | hypothetical protein |
| 2861569 | 2871301 | 9732 | 2869595 | 2871301 | -1 | hypothetical protein |
| 2861569 | 2871301 | 9732 | 2871298 | 2873826 | -1 | VgrG protein |
| 3341988 | 3358673 | 16685 | 3341372 | 3341995 | -1 | Ribosomal-protein-S5p-alanine acetyltransferase (EC 2.3.1.128) |
| 3341988 | 3358673 | 16685 | 3341988 | 3342284 | -1 | hypothetical protein |
| 3341988 | 3358673 | 16685 | 3342593 | 3343888 | 1 | Methyl-accepting chemotaxis sensor/transducer protein |
| 3341988 | 3358673 | 16685 | 3343979 | 3344140 | 1 | hypothetical protein |
| 3341988 | 3358673 | 16685 | 3344305 | 3344685 | 1 | hypothetical protein |
| 3341988 | 3358673 | 16685 | 3345458 | 3346405 | 1 | hypothetical protein |
| 3341988 | 3358673 | 16685 | 3346402 | 3347355 | 1 | Retron-type RNA-directed DNA polymerase (EC 2.7.7.49) |
| 3341988 | 3358673 | 16685 | 3347509 | 3349491 | 1 | hypothetical protein |
| 3341988 | 3358673 | 16685 | 3349488 | 3349610 | 1 | hypothetical protein |
| 3341988 | 3358673 | 16685 | 3349600 | 3351264 | 1 | hypothetical protein |
| 3341988 | 3358673 | 16685 | 3351429 | 3352049 | 1 | Phage integrase |
| 3341988 | 3358673 | 16685 | 3352252 | 3353208 | -1 | hypothetical protein |
| 3341988 | 3358673 | 16685 | 3353579 | 3353719 | 1 | hypothetical protein |
| 3341988 | 3358673 | 16685 | 3353759 | 3354898 | -1 | Catalase-like heme-binding protein |
| 3341988 | 3358673 | 16685 | 3355078 | 3356973 | -1 | hypothetical protein |
| 3341988 | 3358673 | 16685 | 3357284 | 3357418 | -1 | hypothetical protein |
| 3341988 | 3358673 | 16685 | 3357576 | 3358673 | -1 | GNAT family acetyltransferase VC2332 |
| 3343979 | 3349491 | 5512 | 3343979 | 3344140 | 1 | hypothetical protein |
| 3343979 | 3349491 | 5512 | 3344305 | 3344685 | 1 | hypothetical protein |
| 3343979 | 3349491 | 5512 | 3345458 | 3346405 | 1 | hypothetical protein |
| 3343979 | 3349491 | 5512 | 3346402 | 3347355 | 1 | Retron-type RNA-directed DNA polymerase (EC 2.7.7.49) |
| 3343979 | 3349491 | 5512 | 3347509 | 3349491 | 1 | hypothetical protein |
| 3343979 | 3349491 | 5512 | 3349488 | 3349610 | 1 | hypothetical protein |
| 3349600 | 3353719 | 4119 | 3349488 | 3349610 | 1 | hypothetical protein |
| 3349600 | 3353719 | 4119 | 3349600 | 3351264 | 1 | hypothetical protein |
| 3349600 | 3353719 | 4119 | 3351429 | 3352049 | 1 | Phage integrase |
| 3349600 | 3353719 | 4119 | 3352252 | 3353208 | -1 | hypothetical protein |
| 3349600 | 3353719 | 4119 | 3353579 | 3353719 | 1 | hypothetical protein |
| 3545235 | 3549275 | 4040 | 3544294 | 3545250 | 1 | 6-hexanolactone hydrolase |
| 3545235 | 3549275 | 4040 | 3545235 | 3545990 | 1 | Putative short-chain dehydrogenase |
| 3545235 | 3549275 | 4040 | 3546211 | 3547644 | -1 | Transcriptional regulator, GntR family domain / Aspartate aminotransferase (EC 2.6.1.1) |
| 3545235 | 3549275 | 4040 | 3547777 | 3548667 | 1 | Permease of the drug/metabolite transporter (DMT) superfamily |
| 3545235 | 3549275 | 4040 | 3548961 | 3549275 | 1 | hypothetical protein |
| 3660206 | 3671841 | 11635 | 3660206 | 3660322 | 1 | hypothetical protein |
| 3660206 | 3671841 | 11635 | 3660319 | 3661527 | -1 | Putative transcriptional regulator |
| 3660206 | 3671841 | 11635 | 3661743 | 3662654 | 1 | Transcriptional regulator, LysR family |
| 3660206 | 3671841 | 11635 | 3662891 | 3663466 | -1 | hypothetical protein |
| 3660206 | 3671841 | 11635 | 3663698 | 3664339 | -1 | hypothetical protein |
| 3660206 | 3671841 | 11635 | 3664926 | 3665258 | 1 | hypothetical protein |
| 3660206 | 3671841 | 11635 | 3665515 | 3666621 | 1 | hypothetical protein |
| 3660206 | 3671841 | 11635 | 3666831 | 3669746 | 1 | DNA helicase IV (EC 3.6.4.12) |
| 3660206 | 3671841 | 11635 | 3670491 | 3670697 | 1 | Transposase |
| 3660206 | 3671841 | 11635 | 3670801 | 3671841 | 1 | Transposase |
| 3672978 | 3678199 | 5221 | 3672978 | 3673997 | -1 | hypothetical protein |
| 3672978 | 3678199 | 5221 | 3674030 | 3674758 | -1 | hypothetical protein |
| 3672978 | 3678199 | 5221 | 3675007 | 3675423 | -1 | hypothetical protein |
| 3672978 | 3678199 | 5221 | 3676002 | 3676979 | -1 | Transcriptional regulator, AraC family |
| 3672978 | 3678199 | 5221 | 3676988 | 3678199 | -1 | Beta-lactamase class C-like and penicillin binding proteins (PBPs) superfamily |
| 3820853 | 3828405 | 7552 | 3820172 | 3821050 | -1 | MBL-fold metallo-hydrolase superfamily |
| 3820853 | 3828405 | 7552 | 3821112 | 3822437 | -1 | Uncharacterized MFS-type transporter |
| 3820853 | 3828405 | 7552 | 3822691 | 3822816 | -1 | RidA/YER057c/UK114 superfamily protein |
| 3820853 | 3828405 | 7552 | 3823155 | 3824270 | -1 | Pyridoxal-5'-phosphate-dependent enzyme beta superfamily (fold type II) |
| 3820853 | 3828405 | 7552 | 3824392 | 3825636 | -1 | Beta-ureidopropionase (EC 3.5.1.6) |
| 3820853 | 3828405 | 7552 | 3825651 | 3826172 | -1 | L-2-amino-thiazoline-4-carboxylic acid hydrolase (EC 3.5.2.-) |
| 3820853 | 3828405 | 7552 | 3826357 | 3827208 | 1 | hypothetical protein |
| 3820853 | 3828405 | 7552 | 3827284 | 3828027 | -1 | Arylmalonate decarboxylase (EC 4.1.1.76) |
| 3820853 | 3828405 | 7552 | 3828387 | 3829517 | 1 | Glycerophosphoryl diester phosphodiesterase (EC 3.1.4.46) periplasmic (secreted in GramPositives) |
| 3930168 | 3948375 | 18207 | 3930168 | 3930848 | 1 | Nicotinamidase family protein YcaC |
| 3930168 | 3948375 | 18207 | 3930929 | 3932347 | 1 | Outer membrane low permeability porin, OprD family => OccD8/OpdJ coexpressed with pyoverdine biosynt |
| 3930168 | 3948375 | 18207 | 3932380 | 3933300 | 1 | Hypothertical protein, coexpressed with pyoverdine biosynthesis regulon |
| 3930168 | 3948375 | 18207 | 3933431 | 3934447 | 1 | Mobile element protein |
| 3930168 | 3948375 | 18207 | 3935262 | 3935576 | 1 | hypothetical protein |
| 3930168 | 3948375 | 18207 | 3935761 | 3935940 | -1 | hypothetical protein |
| 3930168 | 3948375 | 18207 | 3936228 | 3936425 | -1 | hypothetical protein |
| 3930168 | 3948375 | 18207 | 3936797 | 3937771 | 1 | CRISPR-associated protein Cas1 |
| 3930168 | 3948375 | 18207 | 3937768 | 3940998 | 1 | CRISPR-associated helicase Cas3 |
| 3930168 | 3948375 | 18207 | 3941407 | 3942708 | 1 | CRISPR-associated protein, Csy1 family |
| 3930168 | 3948375 | 18207 | 3942695 | 3943678 | 1 | CRISPR-associated protein, Csy2 family |
| 3930168 | 3948375 | 18207 | 3943689 | 3944717 | 1 | CRISPR-associated protein, Csy3 family |
| 3930168 | 3948375 | 18207 | 3944721 | 3945284 | 1 | CRISPR-associated protein, Csy4 family |
| 3930168 | 3948375 | 18207 | 3945562 | 3946050 | 1 | FIG00613345: hypothetical protein |
| 3930168 | 3948375 | 18207 | 3946450 | 3946710 | 1 | hypothetical protein |
| 3930168 | 3948375 | 18207 | 3946710 | 3947027 | 1 | hypothetical protein |
| 3930168 | 3948375 | 18207 | 3947570 | 3948082 | 1 | Prophage antirepressor |
| 3930168 | 3948375 | 18207 | 3948217 | 3948375 | 1 | hypothetical protein |
| 4139742 | 4144224 | 4482 | 4139742 | 4142627 | -1 | Virulence sensor protein bvgS precursor (EC 2.7.3.-) |
| 4139742 | 4144224 | 4482 | 4143169 | 4144224 | 1 | hypothetical protein |
| 4227461 | 4232529 | 5068 | 4227461 | 4227778 | -1 | hypothetical protein |
| 4227461 | 4232529 | 5068 | 4228021 | 4228605 | -1 | hypothetical protein |
| 4227461 | 4232529 | 5068 | 4228605 | 4229048 | -1 | hypothetical protein |
| 4227461 | 4232529 | 5068 | 4229635 | 4230714 | -1 | Type II secretory pathway, component PulK |
| 4227461 | 4232529 | 5068 | 4230704 | 4231294 | -1 | Type II secretory pathway, pseudopilin PulG |
| 4227461 | 4232529 | 5068 | 4231291 | 4231716 | -1 | General secretion pathway protein I |
| 4227461 | 4232529 | 5068 | 4231716 | 4232126 | -1 | Probable type II secretion system protein |
| 4227461 | 4232529 | 5068 | 4232095 | 4232529 | -1 | General secretion pathway protein G |
| 4288619 | 4312134 | 23515 | 4288619 | 4289098 | 1 | 3-demethylubiquinone-9 3-methyltransferase |
| 4288619 | 4312134 | 23515 | 4289130 | 4289522 | 1 | Gfa-like protein |
| 4288619 | 4312134 | 23515 | 4289552 | 4289830 | 1 | hypothetical protein |
| 4288619 | 4312134 | 23515 | 4289905 | 4290447 | 1 | hypothetical protein |
| 4288619 | 4312134 | 23515 | 4290578 | 4292407 | 1 | Chaperone protein ClpB (ATP-dependent unfoldase) |
| 4288619 | 4312134 | 23515 | 4292409 | 4293047 | 1 | Ribonucleotide reductase of class III (anaerobic), activating protein (EC 1.97.1.4) |
| 4288619 | 4312134 | 23515 | 4293062 | 4299358 | 1 | Superfamily I DNA and RNA helicases and helicase subunits |
| 4288619 | 4312134 | 23515 | 4299355 | 4300704 | 1 | hypothetical protein |
| 4288619 | 4312134 | 23515 | 4301048 | 4301902 | 1 | hypothetical protein |
| 4288619 | 4312134 | 23515 | 4303464 | 4304057 | 1 | Transposase |
| 4288619 | 4312134 | 23515 | 4304079 | 4304264 | 1 | COG2801: Transposase and inactivated derivatives |
| 4288619 | 4312134 | 23515 | 4304442 | 4307630 | 1 | DEAD/DEAH box helicase-like protein |
| 4288619 | 4312134 | 23515 | 4307634 | 4308389 | 1 | Methyl-accepting chemotaxis protein |
| 4288619 | 4312134 | 23515 | 4308524 | 4310551 | 1 | Type III restriction-modification system methylation subunit (EC 2.1.1.72) |
| 4288619 | 4312134 | 23515 | 4310553 | 4310996 | 1 | hypothetical protein |
| 4288619 | 4312134 | 23515 | 4311076 | 4312134 | 1 | ADP-ribose 1-phosphate phophatase related protein |
| 4304442 | 4310996 | 6554 | 4304442 | 4307630 | 1 | DEAD/DEAH box helicase-like protein |
| 4304442 | 4310996 | 6554 | 4307634 | 4308389 | 1 | Methyl-accepting chemotaxis protein |
| 4304442 | 4310996 | 6554 | 4308524 | 4310551 | 1 | Type III restriction-modification system methylation subunit (EC 2.1.1.72) |
| 4304442 | 4310996 | 6554 | 4310553 | 4310996 | 1 | hypothetical protein |
| 4312840 | 4317355 | 4515 | 4312245 | 4315259 | 1 | Type III restriction enzyme, res subunit:DEAD/DEAH box helicase, N-terminal |
| 4312840 | 4317355 | 4515 | 4315729 | 4317630 | -1 | putative DNA helicase |
| 4751735 | 4756139 | 4404 | 4750944 | 4751738 | -1 | Putative N-acetylgalactosaminyl-diphosphoundecaprenol glucuronosyltransferase |
| 4751735 | 4756139 | 4404 | 4751735 | 4753288 | -1 | hypothetical protein |
| 4751735 | 4756139 | 4404 | 4753362 | 4754417 | -1 | N-acetylneuraminate synthase (EC 2.5.1.56) |
| 4751735 | 4756139 | 4404 | 4754417 | 4755067 | -1 | hypothetical protein |
| 4751735 | 4756139 | 4404 | 4755060 | 4756139 | -1 | N-Acetylneuraminate cytidylyltransferase (EC 2.7.7.43) |
| 5458876 | 5472832 | 13956 | 5458876 | 5461398 | 1 | ATP-dependent helicase HrpB |
| 5458876 | 5472832 | 13956 | 5461976 | 5462239 | 1 | Insertion element IS407 (Burkholderia multivorans) transposase |
| 5458876 | 5472832 | 13956 | 5462272 | 5463114 | 1 | Mobile element protein |
| 5458876 | 5472832 | 13956 | 5463662 | 5464270 | -1 | hypothetical protein |
| 5458876 | 5472832 | 13956 | 5464757 | 5466775 | -1 | hypothetical protein |
| 5458876 | 5472832 | 13956 | 5467640 | 5468566 | 1 | hypothetical protein |
| 5458876 | 5472832 | 13956 | 5468845 | 5469654 | -1 | hypothetical protein |
| 5458876 | 5472832 | 13956 | 5469730 | 5469888 | -1 | hypothetical protein |
| 5458876 | 5472832 | 13956 | 5469962 | 5471656 | 1 | ATP-dependent helicase HrpB |
| 5458876 | 5472832 | 13956 | 5471667 | 5472074 | -1 | hypothetical protein |
| 5458876 | 5472832 | 13956 | 5472098 | 5472832 | -1 | ABC-type amino acid transport/signal transduction system, periplasmic component/domain |
| 5462272 | 5469654 | 7382 | 5462272 | 5463114 | 1 | Mobile element protein |
| 5462272 | 5469654 | 7382 | 5463662 | 5464270 | -1 | hypothetical protein |
| 5462272 | 5469654 | 7382 | 5464757 | 5466775 | -1 | hypothetical protein |
| 5462272 | 5469654 | 7382 | 5467640 | 5468566 | 1 | hypothetical protein |
| 5462272 | 5469654 | 7382 | 5468845 | 5469654 | -1 | hypothetical protein |
| 5908254 | 5913950 | 5696 | 5908254 | 5908568 | 1 | hypothetical protein |
| 5908254 | 5913950 | 5696 | 5908753 | 5909805 | -1 | hypothetical protein |
| 5908254 | 5913950 | 5696 | 5910276 | 5910584 | 1 | Transposase InsN for insertion sequence element IS911 |
| 5908254 | 5913950 | 5696 | 5910611 | 5911438 | 1 | Transposase InsO for insertion sequence element IS911 |
| 5908254 | 5913950 | 5696 | 5911901 | 5912323 | -1 | hypothetical protein |
| 5908254 | 5913950 | 5696 | 5912423 | 5912953 | -1 | Cytochrome b561 |
| 5908254 | 5913950 | 5696 | 5913136 | 5913333 | 1 | Copper(I) chaperone CopZ |
| 5908254 | 5913950 | 5696 | 5913312 | 5913950 | -1 | Transcriptional regulator, AcrR family |
| 6072400 | 6077771 | 5371 | 6072400 | 6073416 | -1 | Mobile element protein |
| 6072400 | 6077771 | 5371 | 6073727 | 6074086 | -1 | pyocin R2_PP, conserved hypothetical protein |
| 6072400 | 6077771 | 5371 | 6074134 | 6074334 | -1 | Phage TraR/YbiI family protein |
| 6072400 | 6077771 | 5371 | 6074384 | 6074518 | 1 | hypothetical protein |
| 6072400 | 6077771 | 5371 | 6074791 | 6075561 | 1 | Phage repressor protein cI |
| 6072400 | 6077771 | 5371 | 6075661 | 6075975 | 1 | hypothetical protein |
| 6072400 | 6077771 | 5371 | 6076293 | 6077771 | -1 | Anthranilate synthase, aminase component (EC 4.1.3.27) |

**Table S4. Genomic islands on Iso 1 genome predicted by IslandViewer 4.**
