## Supplementary Table S5 for "Persistent bacterial coinfection of a COVID-19 patient caused by a genetically adapted *Pseudomonas aeruginosa* chronic colonizer"

| **Type** | **Amino acid change** | **Non-synonymous** | **Gene** | **Product** |
| --- | --- | --- | --- | --- |
| Deletion | Pro326_Pro327del | Yes | ppkA | T6SS HSI-1 |
| SNV |  | - |  |  |
| SNV |  | - |  |  |
| Deletion |  | - |  |  |
| SNV | Glu519Asp | Yes | ostA | organic solvent tolerance protein OstA |
| Insertion | Arg111_Arg112insLeuArg | Yes | PA0700 | hypothetical protein |
| SNV |  | - |  |  |
| SNV |  | - |  |  |
| Insertion | Glu171fs | Yes | oprD | porin D |
| SNV |  | No | fgtA | flagellar glycosyl transferase FgtA |
| Deletion |  | - |  |  |
| Deletion | Trp463fs | Yes | PA1194 | amino acid permease |
| Deletion | Gly206_Asp207del | Yes | PA1359 | transcriptional regulator |
| SNV |  | No | PA1658 | HsiC2 (T6SS HSI-2) |
| SNV |  | No | PA1658 | HsiC2 (T6SS HSI-2) |
| SNV |  | No | PA1658 | HsiC2 (T6SS HSI-2) |
| SNV |  | No | PA1658 | HsiC2 (T6SS HSI-2) |
| SNV |  | No | PA1658 | HsiC2 (T6SS HSI-2) |
| SNV |  | No | PA1658 | HsiC2 (T6SS HSI-2) |
| SNV |  | No | PA1938 | hypothetical protein |
| Deletion |  | - |  |  |
| MNV |  | - |  |  |
| SNV |  | - |  |  |
| SNV |  | - |  |  |
| SNV |  | - |  |  |
| SNV |  | - |  |  |
| SNV |  | No | pvdA | L-ornithine N5-oxygenase |
| SNV |  | No | pvdA | L-ornithine N5-oxygenase |
| MNV | Ile210Val | Yes | pvdA | L-ornithine N5-oxygenase |
| SNV |  | No | pvdA | L-ornithine N5-oxygenase |
| SNV |  | No | pvdP | pyoverdine biosynthesis protein PvdP |
| MNV | Val114Leu | Yes | pvdP | pyoverdine biosynthesis protein PvdP |
| SNV |  | No | pvdP | pyoverdine biosynthesis protein PvdP |
| MNV | Thr31Val | Yes | pvdP | pyoverdine biosynthesis protein PvdP |
| SNV | Ser1348Cys | Yes | pvdD | pyoverdine synthetase D |
| Insertion | Ser1347fs | Yes | pvdD | pyoverdine synthetase D |
| SNV |  | No | pvdD | pyoverdine synthetase D |
| MNV | Ala1346Thr | Yes | pvdD | pyoverdine synthetase D |
| SNV |  | No | pvdD | pyoverdine synthetase D |
| SNV |  | No | pvdD | pyoverdine synthetase D |
| SNV |  | No | pvdD | pyoverdine synthetase D |
| SNV |  | No | pvdD | pyoverdine synthetase D |
| SNV |  | No | PA2402 | peptide synthase |
| SNV |  | No | PA2402 | peptide synthase |
| SNV | Glu4765Ala | Yes | PA2402 | peptide synthase |
| SNV |  | No | PA2402 | peptide synthase |
| SNV |  | No | PA2402 | peptide synthase |
| SNV |  | No | PA2402 | peptide synthase |
| SNV |  | No | PA2402 | peptide synthase |
| SNV |  | No | PA2402 | peptide synthase |
| MNV | Cys1794Thr | Yes | PA2402 | peptide synthase |
| MNV | His1793Tyr | Yes | PA2402 | peptide synthase |
| SNV |  | No | PA2402 | peptide synthase |
| SNV |  | No | PA2402 | peptide synthase |
| SNV |  | No | PA2402 | peptide synthase |
| SNV | Ala1787Ser | Yes | PA2402 | peptide synthase |
| SNV |  | No | pvdL | peptide synthase |
| SNV |  | No | PA2438 | hypothetical protein |
| SNV |  | - |  |  |
| Deletion |  | - |  |  |
| Insertion | Ala148_Gly149insValAla | Yes | PA2478 | thiol:disulfide interchange protein DsbD |
| Deletion |  | - |  |  |
| Deletion | Leu8fs | Yes | PA3053 | hydrolase |
| SNV |  | - |  |  |
| MNV |  | - |  |  |
| SNV |  | - |  |  |
| Insertion |  | - |  |  |
| SNV |  | - |  |  |
| MNV |  | - |  |  |
| SNV |  | - |  |  |
| MNV | Pro634Gly | Yes | xcpQ | type II secretion system protein D |
| SNV |  | No | xcpQ | type II secretion system protein D |
| SNV |  | - |  |  |
| Insertion |  | - |  |  |
| SNV |  | - |  |  |
| Insertion |  | - |  |  |
| SNV |  | - |  |  |
| Replacement |  | - |  |  |
| SNV |  | - |  |  |
| Insertion |  | - |  |  |
| SNV |  | No | PA3294 | VgrG4a |
| Insertion |  | - |  |  |
| Deletion | Gln231_Arg232del | Yes | PA3494 | VgrG4a |
| Insertion |  | - |  |  |
| Replacement |  | - |  |  |
| SNV |  | - |  |  |
| SNV |  | - |  |  |
| SNV |  | - |  |  |
| SNV |  | - |  |  |
| SNV |  | - |  |  |
| Insertion |  | - |  |  |
| Insertion |  | - |  |  |
| Insertion |  | - |  |  |
| Insertion |  | - |  |  |
| Insertion |  | - |  |  |
| SNV |  | - |  |  |
| SNV |  | - |  |  |
| SNV |  | - |  |  |
| MNV |  | - |  |  |
| Insertion |  | - |  |  |
| Deletion | Gln484del | Yes | PA4072 | amino acid permease |
| Insertion |  | - |  |  |
| Insertion |  | - |  |  |
| Insertion |  | - |  |  |
| Deletion |  | - |  |  |
| Insertion |  | - |  |  |
| SNV |  | No | pctB | chemotactic transducer PctB |
| Insertion |  | - |  |  |
| SNV |  | - |  |  |
| Insertion |  | - |  |  |
| Replacement |  | - |  |  |
| SNV |  | - |  |  |
| Deletion |  | - |  |  |
| SNV |  | No | PA4625 | hypothetical protein |
| Deletion |  | - |  |  |
| Insertion | .Leu100dup | Yes | PA4936 | 23S rRNA (guanosine(2251)-2'-O)-methyltransferase RlmB |
| SNV |  | No | PA5284 | hypothetical protein |
| SNV |  | No | PA5284 | hypothetical protein |

**Table S5. List of SNPs of Iso 2 comparing to Iso 1.**
