## Supplementary Table S6 for "Persistent bacterial coinfection of a COVID-19 patient caused by a genetically adapted *Pseudomonas aeruginosa* chronic colonizer"

| **Feature ID** | **Product Name** | **Fold change** | **P-value** | **Adjusted p-value** | **Iso 1 Means** | **Iso 2 Means** |
| --- | --- | --- | --- | --- | --- | --- |
| acoR | transcriptional regulator AcoR | -44.08 | 5.39E-06 | 2.08E-03 | 16.5 | 0.5 |
| exoS | exoenzyme S | -73.60 | 1.64E-05 | 3.99E-03 | 7.5 | 0 |
| fpvA | ferripyoverdine receptor | -11.93 | 1.82E-04 | 2.20E-02 | 55.5 | 8 |
| glpD | glycerol-3-phosphate dehydrogenase | -4.14 | 2.82E-14 | 4.02E-11 | 1805.5 | 674 |
| mexC | Resistance-Nodulation-Cell Division (RND) multidrug efflux membrane fusion protein MexC precursor | -8.22 | 4.80E-04 | 4.27E-02 | 2709.5 | 583 |
| mexD | Resistance-Nodulation-Cell Division (RND) multidrug efflux transporter MexD | -12.15 | 2.03E-04 | 2.41E-02 | 5469 | 802.5 |
| PA0053 | hypothetical protein | -249.24 | 1.38E-08 | 1.12E-05 | 23 | 0 |
| PA0257 | hypothetical protein | -213.99 | 2.46E-05 | 5.20E-03 | 19 | 0 |
| PA0259 | Type 6 lipase adaptor, Tla3 | -97.73 | 1.50E-05 | 3.87E-03 | 36 | 0.5 |
| PA0260 | Type 6 lipase effector, Tle3 | -32.38 | 1.94E-05 | 4.42E-03 | 30 | 1.5 |
| PA0261 | Type 6 lipase immunity, Tli3 | -17.53 | 1.40E-04 | 2.00E-02 | 7 | 0.5 |
| PA0497 | hypothetical protein | -4.23 | 1.03E-04 | 1.54E-02 | 26.5 | 10 |
| PA0561 | hypothetical protein | -228.36 | 2.34E-06 | 1.02E-03 | 20.5 | 0 |
| PA0689 | low-molecular-weight alkaline phosphatase B, LapB | -51.63 | 9.51E-06 | 3.01E-03 | 19 | 0.5 |
| PA0701a |  | -55.40 | 5.59E-04 | 4.72E-02 | 5.5 | 0 |
| PA1194 | probable amino acid permease | -68.25 | 3.53E-04 | 3.66E-02 | 6.5 | 0 |
| PA1241 | probable transcriptional regulator | -248.95 | 4.26E-05 | 7.83E-03 | 22 | 0 |
| PA1312 | probable transcriptional regulator | -61.40 | 2.68E-04 | 2.94E-02 | 6 | 0 |
| PA1315 | probable transcriptional regulator | -85.79 | 1.45E-04 | 2.02E-02 | 8 | 0 |
| PA1428a |  | -100.82 | 1.34E-05 | 3.81E-03 | 9.5 | 0 |
| PA1922 | probable TonB-dependent receptor | -4.02 | 4.56E-04 | 4.26E-02 | 92 | 38 |
| PA1933 | probable hydroxylase large subunit | -14.01 | 1.66E-04 | 2.19E-02 | 6 | 0.5 |
| PA2037 | hypothetical protein | -150.60 | 5.12E-05 | 8.84E-03 | 13.5 | 0 |
| PA2119 | alcohol dehydrogenase (Zn-dependent) | -69.74 | 6.41E-06 | 2.22E-03 | 44.5 | 1 |
| PA2459 | hypothetical protein | -83.88 | 7.75E-05 | 1.19E-02 | 8 | 0 |
| PA2564 | hypothetical protein | -55.64 | 6.13E-04 | 4.99E-02 | 5.5 | 0 |
| PA2594 | conserved hypothetical protein | -40.02 | 1.21E-05 | 3.63E-03 | 15 | 0.5 |
| PA2601 | probable transcriptional regulator | -89.72 | 3.85E-05 | 7.32E-03 | 8.5 | 0 |
| PA2603 | probable thiosulfate sulfurtransferase | -54.56 | 4.11E-04 | 4.05E-02 | 5.5 | 0 |
| PA2764 | hypothetical protein | -155.49 | 1.68E-05 | 3.99E-03 | 14 | 0 |
| PA2771 | diguanylate cyclase with a self-blocked I-site, Dcsbis | -39.88 | 5.48E-06 | 2.08E-03 | 15 | 0.5 |
| PA2775 | Tsi4 | -127.78 | 1.19E-04 | 1.73E-02 | 11.5 | 0 |
| PA3362 | hypothetical protein | -59.62 | 2.28E-04 | 2.55E-02 | 6 | 0 |
| PA4191 | isopenicillin-N synthase | -89.64 | 3.70E-05 | 7.26E-03 | 8.5 | 0 |
| PA4802 | hypothetical protein | -67.99 | 3.07E-04 | 3.30E-02 | 6.5 | 0 |
| PA4849 | hypothetical protein | -174.20 | 7.43E-05 | 1.18E-02 | 15.5 | 0 |
| PA5264 | hypothetical protein | -297.40 | 1.35E-06 | 7.01E-04 | 26.5 | 0 |
| PA5265 | hypothetical protein | -108.26 | 6.62E-06 | 2.22E-03 | 40 | 0.5 |
| pys2 | pyocin S2 | -16.90 | 1.79E-04 | 2.20E-02 | 34.5 | 3.5 |

**Table S6. Significantly regulated genes in Iso 2 comparing to that of Iso 1. (Absolute fold change ≧ 4, adjusted p-value < 0.05)**
