## Supplementary Table S7 for "Persistent bacterial coinfection of a COVID-19 patient caused by a genetically adapted *Pseudomonas aeruginosa* chronic colonizer"

| **Feature ID** | **Product** | **Fold change** | **P-value** | **Adjusted p-value** | **PAO1 Means** | **Iso 2 Means** |
| --- | --- | --- | --- | --- | --- | --- |
| exaC | NAD+ dependent aldehyde dehydrogenase ExaC | 68.60 | 1.48E-85 | 6.45E-84 | 20 | 1980 |
| PA1541 | probable drug efflux transporter | 66.23 | 2.54E-101 | 1.28E-99 | 6 | 579 |
| PA0978 | conserved hypothetical protein | 40.58 | 3.65E-91 | 1.72E-89 | 20.5 | 1199 |
| norC | nitric-oxide reductase subunit C | 40.00 | 6.93E-103 | 3.56E-101 | 18 | 1032 |
| PA2019 | Resistance-Nodulation-Cell Division (RND) multidrug efflux membrane fusion protein MexX precursor | 38.30 | 2.11E-81 | 8.90E-80 | 78 | 4281.5 |
| PA2421 | hypothetical protein | 35.16 | 1.28E-77 | 5.18E-76 | 7.5 | 383.5 |
| PA0979 | conserved hypothetical protein | 28.30 | 1.45E-118 | 8.24E-117 | 23.5 | 955.5 |
| PA2018 | Resistance-Nodulation-Cell Division (RND) multidrug efflux transporter MexY | 28.23 | 3.50E-93 | 1.68E-91 | 353 | 14283 |
| PA0986 | conserved hypothetical protein | 28.05 | 3.85E-45 | 1.13E-43 | 5 | 205.5 |
| nosR | regulatory protein NosR | 27.74 | 1.09E-105 | 5.80E-104 | 59.5 | 2359 |
| PA3661 | hypothetical protein | 26.66 | 2.06E-66 | 7.14E-65 | 9.5 | 368 |
| PA1781.1 | nitrogen assimilation leader A (NalA) | 26.30 | 1.90E-02 | 3.76E-02 | 0 | 4 |
| norB | nitric-oxide reductase subunit B | 22.20 | 1.69E-86 | 7.47E-85 | 71.5 | 2260.5 |
| algA | phosphomannose isomerase / guanosine 5'-diphospho-D-mannose pyrophosphorylase | 20.08 | 2.11E-65 | 7.30E-64 | 39 | 1132 |
| PA2422 | hypothetical protein | 19.58 | 2.01E-37 | 5.31E-36 | 5 | 143 |
| PA2107 | hypothetical protein | 18.56 | 5.45E-68 | 1.90E-66 | 15 | 402.5 |
| nosZ | nitrous-oxide reductase precursor | 16.97 | 1.31E-104 | 6.84E-103 | 54.5 | 1324 |
| PA0987 | conserved hypothetical protein | 16.29 | 1.15E-78 | 4.80E-77 | 24.5 | 574.5 |
| algD | GDP-mannose 6-dehydrogenase AlgD | 15.91 | 3.78E-55 | 1.19E-53 | 45.5 | 1049.5 |
| kdpC | potassium-transporting ATPase, C chain | 15.59 | 3.28E-18 | 4.84E-17 | 8 | 181.5 |
| algF | alginate o-acetyltransferase AlgF | 13.99 | 3.22E-47 | 9.57E-46 | 14.5 | 294.5 |
| algX | alginate biosynthesis protein AlgX | 13.82 | 2.54E-47 | 7.56E-46 | 13 | 260.5 |
| fhp | flavohemoprotein | 13.45 | 1.70E-81 | 7.25E-80 | 58 | 1115.5 |
| PA4139 | hypothetical protein | 12.91 | 2.46E-40 | 6.78E-39 | 107 | 1996.5 |
| PA0982 | hypothetical protein | 12.40 | 1.19E-99 | 5.90E-98 | 104.5 | 1852.5 |
| qscR | quorum-sensing control repressor | 11.99 | 6.68E-90 | 3.07E-88 | 77.5 | 1330 |
| PA4173 | conserved hypothetical protein | 11.89 | 4.72E-20 | 7.74E-19 | 5.5 | 95.5 |
| algE | Alginate production outer membrane protein AlgE precursor | 11.06 | 8.52E-52 | 2.65E-50 | 21.5 | 343 |
| czcC | outer membrane protein precursor CzcC | 10.53 | 7.27E-29 | 1.62E-27 | 11.5 | 174 |
| algJ | alginate o-acetyltransferase AlgJ | 10.34 | 6.24E-26 | 1.29E-24 | 9.5 | 142.5 |
| algL | poly(beta-d-mannuronate) lyase precursor AlgL | 10.08 | 1.59E-46 | 4.68E-45 | 19 | 277 |
| PA2171 | hypothetical protein | 9.74 | 9.25E-20 | 1.50E-18 | 10.5 | 148.5 |
| PA4582 | conserved hypothetical protein | 9.51 | 2.05E-72 | 7.64E-71 | 90 | 1231.5 |
| kdpB | potassium-transporting ATPase, B chain | 9.44 | 5.89E-50 | 1.80E-48 | 49.5 | 672 |
| PA0087 | TssE1 | 9.27 | 6.57E-72 | 2.38E-70 | 95.5 | 1269.5 |
| PA1896 | hypothetical protein | 9.24 | 1.67E-87 | 7.50E-86 | 336.5 | 4461 |
| PA0320 | calcium-regulated OB-fold protein CarO | 8.93 | 2.53E-70 | 8.95E-69 | 49 | 629 |
| PA0086 | TagJ1 | 8.89 | 3.21E-70 | 1.13E-68 | 416 | 5304 |
| PA4142 | probable secretion protein | 8.87 | 2.27E-71 | 8.15E-70 | 97 | 1227.5 |
| PA4140 | hypothetical protein | 8.85 | 2.87E-44 | 8.23E-43 | 70 | 894 |
| algK | alginate biosynthetic protein AlgK precursor | 8.80 | 2.57E-26 | 5.39E-25 | 15.5 | 197.5 |
| PA1891 | hypothetical protein | 8.72 | 5.74E-61 | 1.92E-59 | 30 | 376 |
| PA0534 | FAD-dependent oxidoreductase | 8.69 | 3.29E-73 | 1.24E-71 | 202 | 2513.5 |
| PA1844 | Tse1 | 8.53 | 1.34E-38 | 3.58E-37 | 42 | 516.5 |
| PA2497 | probable transcriptional regulator | 8.46 | 1.94E-82 | 8.29E-81 | 74.5 | 902 |
| alg44 | alginate biosynthesis protein Alg44 | 8.27 | 7.19E-33 | 1.79E-31 | 22 | 263 |
| nosD | NosD protein | 7.99 | 3.65E-60 | 1.21E-58 | 35 | 400.5 |
| PA2169 | hypothetical protein | 7.95 | 2.51E-14 | 2.78E-13 | 6.5 | 75 |
| PA1897 | hypothetical protein | 7.80 | 1.10E-37 | 2.91E-36 | 387 | 4344.5 |
| PA3733a |  | 7.50 | 8.66E-58 | 2.79E-56 | 44.5 | 480.5 |
| PA2154 | conserved hypothetical protein | 7.10 | 2.10E-08 | 1.28E-07 | 3.5 | 36.5 |
| PA2319 | probable transposase | 7.05 | 1.11E-03 | 2.98E-03 | 1 | 11 |
| PA1895 | hypothetical protein | 7.02 | 3.47E-73 | 1.30E-71 | 374 | 3763 |
| vgrG1 | VgrG1 | 6.99 | 3.05E-49 | 9.19E-48 | 709.5 | 7118 |
| hcp1 | Hcp1 | 6.91 | 1.92E-71 | 6.93E-70 | 2846 | 28258.5 |
| PA2355 | probable FMNH2-dependent monooxygenase | 6.84 | 5.81E-15 | 6.81E-14 | 16.5 | 162 |
| PA1137 | probable oxidoreductase | 6.71 | 1.63E-30 | 3.83E-29 | 377 | 3608 |
| PA4144 | probable outer membrane protein precursor | 6.69 | 5.15E-58 | 1.67E-56 | 54 | 519 |
| PA1253 | alpha-ketoglutaric semialdehyde dehydrogenase, LhpG | 6.59 | 4.43E-38 | 1.18E-36 | 36.5 | 345 |
| PA2569 | hypothetical protein | 6.55 | 9.20E-20 | 1.49E-18 | 24 | 225 |
| PA2324 | hypothetical protein | 6.48 | 3.81E-24 | 7.44E-23 | 25 | 231.5 |
| PA2496 | hypothetical protein | 6.42 | 4.98E-44 | 1.41E-42 | 66.5 | 612 |
| PA2020 | MexZ | 6.35 | 4.68E-72 | 1.72E-70 | 159 | 1445 |
| PA0525 | probable dinitrification protein NorD | 6.32 | 5.89E-36 | 1.54E-34 | 250 | 2251 |
| PA2036 | hypothetical protein | 6.31 | 2.31E-15 | 2.80E-14 | 11 | 100 |
| PA4143 | probable toxin transporter | 6.30 | 1.04E-73 | 3.96E-72 | 114.5 | 1032.5 |
| PA0522 | hypothetical protein | 6.21 | 5.53E-07 | 2.74E-06 | 3.5 | 32 |
| PA2205 | hypothetical protein | 6.20 | 4.49E-22 | 8.15E-21 | 34 | 302.5 |
| alg8 | alginate biosynthesis protein Alg8 | 6.17 | 4.59E-39 | 1.23E-37 | 43 | 381.5 |
| clpV1 | ClpV1 | 6.08 | 1.08E-44 | 3.10E-43 | 1599 | 13961.5 |
| pelD | PelD | 5.96 | 2.04E-39 | 5.54E-38 | 86.5 | 738 |
| PA2175 | hypothetical protein | 5.95 | 4.35E-10 | 3.24E-09 | 14 | 119 |
| PA1845 | Tsi1 | 5.90 | 1.51E-18 | 2.28E-17 | 39 | 332 |
| PA4583 | conserved hypothetical protein | 5.87 | 2.94E-45 | 8.63E-44 | 161.5 | 1362.5 |
| PA1955 | FapB | 5.80 | 3.85E-05 | 1.42E-04 | 2.5 | 21.5 |
| PA1559 | hypothetical protein | 5.77 | 5.93E-37 | 1.56E-35 | 328.5 | 2716 |
| PA0089 | TssG1 | 5.76 | 2.50E-48 | 7.51E-47 | 227 | 1873 |
| pqqD | pyrroloquinoline quinone biosynthesis protein D | 5.72 | 1.24E-09 | 8.71E-09 | 12 | 99.5 |
| mdcA | malonate decarboxylase alpha subunit | 5.69 | 4.14E-45 | 1.20E-43 | 142 | 1161 |
| PA2221 | conserved hypothetical protein | 5.67 | 2.86E-26 | 5.97E-25 | 31 | 252.5 |
| msuD | methanesulfonate sulfonatase MsuD | 5.66 | 1.19E-09 | 8.37E-09 | 6.5 | 53.5 |
| PA2499 | probable deaminase | 5.63 | 1.19E-15 | 1.47E-14 | 12.5 | 101.5 |
| pelF | PelF | 5.55 | 1.27E-39 | 3.47E-38 | 72 | 573 |
| PA0774 | conserved hypothetical protein | 5.51 | 3.16E-29 | 7.14E-28 | 275 | 2169.5 |
| PA1416 | conserved hypothetical protein | 5.51 | 1.69E-58 | 5.55E-57 | 63.5 | 501 |
| PA2702 | Tse2 | 5.50 | 3.30E-43 | 9.27E-42 | 73 | 575.5 |
| PA1395 | hypothetical protein | 5.44 | 5.62E-21 | 9.71E-20 | 29 | 227.5 |
| PA0445 | probable transposase | 5.37 | 7.37E-05 | 2.57E-04 | 4 | 31.5 |
| PA4624 | cyclic diguanylate-regulated TPS partner B, CdrB | 5.35 | 7.98E-28 | 1.72E-26 | 348.5 | 2681.5 |
| nirL | heme d1 biosynthesis protein NirL | 5.21 | 6.14E-13 | 5.81E-12 | 24.5 | 183 |
| PA0088 | TssF1 | 5.20 | 1.05E-34 | 2.72E-33 | 419.5 | 3128.5 |
| PA1169 | probable lipoxygenase | 5.17 | 1.27E-31 | 3.05E-30 | 151.5 | 1126.5 |
| PA0082 | TssA1 | 5.13 | 5.45E-45 | 1.58E-43 | 237.5 | 1745.5 |
| PA2793 | hypothetical protein | 5.13 | 1.73E-32 | 4.28E-31 | 238 | 1752.5 |
| PA0714 | hypothetical protein | 5.12 | 9.03E-14 | 9.32E-13 | 22 | 161 |
| PA1893 | hypothetical protein | 5.10 | 4.44E-34 | 1.14E-32 | 484 | 3538 |
| PA0512 | NirH | 5.09 | 1.97E-16 | 2.63E-15 | 20 | 146 |
| PA2168 | hypothetical protein | 5.07 | 2.25E-07 | 1.19E-06 | 6.5 | 48 |
| PA2261 | probable 2-ketogluconate kinase | 5.06 | 2.84E-33 | 7.16E-32 | 68.5 | 496.5 |
| PA1894 | hypothetical protein | 5.02 | 6.12E-35 | 1.58E-33 | 513 | 3688.5 |
| PA4584 | conserved hypothetical protein | 4.96 | 1.64E-31 | 3.92E-30 | 106 | 757 |
| PA1977 | hypothetical protein | 4.93 | 7.01E-16 | 8.87E-15 | 20.5 | 145.5 |
| PA2143 | hypothetical protein | 4.93 | 6.83E-08 | 3.92E-07 | 7 | 50 |
| PA0845 | CerN | 4.92 | 1.29E-28 | 2.84E-27 | 90.5 | 635 |
| PA1418 | probable sodium:solute symport protein | 4.90 | 5.53E-19 | 8.49E-18 | 27 | 190 |
| modA | molybdate-binding periplasmic protein precursor ModA | 4.88 | 1.27E-21 | 2.23E-20 | 51 | 356.5 |
| pvcA | paerucumarin biosynthesis protein PvcA | 4.87 | 1.12E-07 | 6.22E-07 | 6.5 | 46 |
| msuE | NADH-dependent FMN reductase MsuE | 4.87 | 4.91E-04 | 1.42E-03 | 2.5 | 18 |
| PA2176 | hypothetical protein | 4.86 | 8.16E-22 | 1.46E-20 | 43 | 300.5 |
| PA2263 | probable 2-hydroxyacid dehydrogenase | 4.86 | 4.46E-19 | 6.94E-18 | 37 | 257.5 |
| PA2260 | hypothetical protein | 4.84 | 1.75E-30 | 4.10E-29 | 68 | 471 |
| PA1415 | hypothetical protein | 4.82 | 6.25E-29 | 1.40E-27 | 341 | 2345.5 |
| PA2134 | hypothetical protein | 4.79 | 1.23E-05 | 4.96E-05 | 4 | 28 |
| PA2792 | hypothetical protein | 4.79 | 4.91E-33 | 1.23E-31 | 123 | 844 |
| PA2703 | Tsi2 | 4.77 | 1.50E-20 | 2.52E-19 | 32 | 219.5 |
| PA1864 | probable transcriptional regulator | 4.75 | 4.82E-12 | 4.25E-11 | 16 | 109.5 |
| pvcD | paerucumarin biosynthesis protein PvcD | 4.73 | 6.14E-06 | 2.59E-05 | 5.5 | 38 |
| PA0072 | TagS1 | 4.70 | 2.13E-29 | 4.83E-28 | 161 | 1082.5 |
| PA1168 | hypothetical protein | 4.69 | 5.88E-22 | 1.06E-20 | 60 | 406.5 |
| nirC | probable c-type cytochrome precursor | 4.68 | 1.03E-21 | 1.82E-20 | 36.5 | 244.5 |
| nirF | heme d1 biosynthesis protein NirF | 4.65 | 5.43E-31 | 1.29E-29 | 159.5 | 1059 |
| PA2125 | probable aldehyde dehydrogenase | 4.63 | 8.14E-30 | 1.88E-28 | 113.5 | 752 |
| PA2667 | MvaU | 4.61 | 1.67E-31 | 3.98E-30 | 823 | 5442.5 |
| PA2314 | probable major facilitator superfamily (MFS) transporter | 4.58 | 4.30E-08 | 2.54E-07 | 8.5 | 56.5 |
| PA1892 | hypothetical protein | 4.56 | 2.00E-33 | 5.08E-32 | 172.5 | 1126.5 |
| PA0097 | hypothetical protein | 4.49 | 3.55E-26 | 7.38E-25 | 174 | 1121.5 |
| pelE | PelE | 4.48 | 5.84E-20 | 9.53E-19 | 51.5 | 329.5 |
| PA0989 | hypothetical protein | 4.43 | 9.88E-26 | 2.03E-24 | 48 | 305 |
| PA0711 | hypothetical protein | 4.39 | 2.87E-10 | 2.18E-09 | 13.5 | 85.5 |
| PA4625 | cyclic diguanylate-regulated TPS partner A, CdrA | 4.37 | 4.24E-18 | 6.23E-17 | 956.5 | 6024.5 |
| PA1540 | conserved hypothetical protein | 4.36 | 1.93E-12 | 1.76E-11 | 20 | 125 |
| PA2146 | conserved hypothetical protein | 4.35 | 5.82E-03 | 1.32E-02 | 2 | 13 |
| PA0084 | TssC1 | 4.35 | 1.02E-18 | 1.54E-17 | 2951.5 | 18477 |
| PA4095 | hypothetical protein | 4.34 | 1.01E-06 | 4.83E-06 | 7 | 44 |
| PA4985 | Uncharacterized protein | 4.30 | 2.28E-19 | 3.64E-18 | 41 | 253 |
| pelG | PelG | 4.29 | 3.77E-19 | 5.93E-18 | 40.5 | 248.5 |
| dctA | C4-dicarboxylate transport protein | 4.29 | 2.70E-27 | 5.79E-26 | 254 | 1564.5 |
| gloA2 | lactoylglutathione lyase | 4.21 | 4.41E-06 | 1.91E-05 | 8.5 | 51.5 |
| PA0515 | probable transcriptional regulator | 4.18 | 1.58E-17 | 2.26E-16 | 38 | 227 |
| PA2160 | probable glycosyl hydrolase | 4.17 | 8.32E-17 | 1.15E-15 | 28.5 | 171 |
| PA5470 | probable peptide chain release factor | 4.16 | 9.55E-24 | 1.82E-22 | 126.5 | 754 |
| PA1400 | probable pyruvate carboxylase | 4.15 | 1.12E-28 | 2.47E-27 | 112 | 666 |
| opdE | membrane protein OpdE | 4.14 | 6.65E-14 | 7.06E-13 | 23 | 136.5 |
| PA0346 | hypothetical protein | 4.11 | 1.66E-16 | 2.25E-15 | 31 | 183 |
| PA2163 | hypothetical protein | 4.11 | 5.47E-12 | 4.80E-11 | 17.5 | 103.5 |
| armR | antirepressor for MexR, ArmR | 4.10 | 2.05E-04 | 6.41E-04 | 4 | 24 |
| PA2415 | hypothetical protein | 4.08 | 1.99E-08 | 1.22E-07 | 11.5 | 68 |
| PA2791 | hypothetical protein | 4.08 | 2.31E-13 | 2.28E-12 | 24.5 | 144 |
| PA0083 | TssB1 | 4.08 | 1.45E-15 | 1.78E-14 | 993 | 5839.5 |
| pqqB | pyrroloquinoline quinone biosynthesis protein B | 4.07 | 2.01E-24 | 3.96E-23 | 108.5 | 635.5 |
| PA2498 | conserved hypothetical protein | 4.06 | 7.11E-11 | 5.76E-10 | 21 | 123 |
| nosF | NosF protein | 4.05 | 1.45E-09 | 1.01E-08 | 19 | 110 |
| PA2218 | hypothetical protein | 4.05 | 1.56E-18 | 2.34E-17 | 71.5 | 415 |
| PA0073 | TagT1 | 4.05 | 6.59E-25 | 1.33E-23 | 186 | 1077 |
| PA2021 | hypothetical protein | 4.01 | 7.82E-09 | 4.98E-08 | 13 | 75 |
| PA2704 | probable transcriptional regulator | 4.01 | 7.36E-19 | 1.12E-17 | 83 | 478.5 |
| PA4691 | hypothetical protein | 4.01 | 4.98E-25 | 1.01E-23 | 79.5 | 457 |
| PA0521 | probable cytochrome c oxidase subunit | 4.00 | 2.56E-11 | 2.13E-10 | 24 | 137.5 |
| PA4714 | conserved hypothetical protein | -4.08 | 0.00E+00 | 0.00E+00 | 1779.5 | 626 |
| PA0165 | hypothetical protein | -4.11 | 3.03E-22 | 5.59E-21 | 297.5 | 103.5 |
| PA2788 | probable chemotaxis transducer | -4.14 | 0.00E+00 | 0.00E+00 | 3152 | 1088.5 |
| PA3054 | hypothetical protein | -4.16 | 8.75E-32 | 2.12E-30 | 1016 | 349 |
| PA0983 | conserved hypothetical protein | -4.19 | 1.90E-09 | 1.30E-08 | 66 | 22.5 |
| grpE | heat shock protein GrpE | -4.24 | 1.15E-14 | 1.30E-13 | 1719 | 583 |
| PA0399 | cystathionine beta-synthase | -4.25 | 0.00E+00 | 0.00E+00 | 4028.5 | 1364 |
| PA3928 | hypothetical protein | -4.29 | 8.92E-07 | 4.31E-06 | 36.5 | 12 |
| PA4933 | hypothetical protein | -4.30 | 0.00E+00 | 0.00E+00 | 26941 | 9018 |
| PA3365 | probable chaperone | -4.31 | 7.22E-27 | 1.53E-25 | 943.5 | 313.5 |
| PA4645 | probable purine/pyrimidine phosphoribosyl transferase | -4.33 | 1.90E-24 | 3.76E-23 | 355 | 117 |
| sahH | S-adenosyl-L-homocysteine hydrolase | -4.35 | 0.00E+00 | 0.00E+00 | 11547.5 | 3813.5 |
| tsf | elongation factor Ts | -4.35 | 0.00E+00 | 0.00E+00 | 21812.5 | 7202 |
| pctC | chemotactic transducer PctC | -4.40 | 3.44E-15 | 4.12E-14 | 1134 | 369 |
| rsmZ | regulatory RNA RsmZ | -4.44 | 6.67E-03 | 1.49E-02 | 75 | 23.5 |
| amiE | aliphatic amidase | -4.56 | 0.00E+00 | 0.00E+00 | 2822.5 | 892.5 |
| dnaK | DnaK protein | -4.59 | 2.22E-16 | 2.95E-15 | 22416 | 7029 |
| PA3291 | Tli1 | -4.67 | 2.86E-14 | 3.15E-13 | 161 | 49.5 |
| PA0605 | AgtC | -4.69 | 1.89E-03 | 4.82E-03 | 1038 | 312.5 |
| PA3284 | hypothetical protein | -4.71 | 1.24E-14 | 1.41E-13 | 178.5 | 54 |
| hslU | heat shock protein HslU | -4.72 | 0.00E+00 | 0.00E+00 | 3720 | 1135.5 |
| gabP |  | -4.76 | 9.97E-21 | 1.68E-19 | 1410 | 425 |
| PA0492 | conserved hypothetical protein | -4.77 | 1.04E-04 | 3.48E-04 | 342.5 | 101.5 |
| PA3486 | VgrG4b | -4.85 | 2.39E-30 | 5.58E-29 | 786.5 | 232 |
| fpr |  | -4.86 | 0.00E+00 | 0.00E+00 | 3484 | 1025 |
| groES | GroES protein | -4.86 | 0.00E+00 | 0.00E+00 | 5445.5 | 1608 |
| glcD | glycolate oxidase subunit GlcD | -4.94 | 4.89E-21 | 8.49E-20 | 1025 | 299 |
| PA0262 | VgrG2b | -5.01 | 2.32E-43 | 6.55E-42 | 745 | 212.5 |
| cioB | cyanide insensitive terminal oxidase | -5.02 | 1.22E-30 | 2.89E-29 | 1458 | 417 |
| PA0779 | AsrA | -5.12 | 0.00E+00 | 0.00E+00 | 4219 | 1182 |
| pchB | salicylate biosynthesis protein PchB | -5.19 | 3.20E-06 | 1.42E-05 | 72 | 19.5 |
| PA0604 | AgtB | -5.20 | 1.52E-08 | 9.43E-08 | 4140.5 | 1131.5 |
| nadE | NH3-dependent NAD synthetase | -5.26 | 0.00E+00 | 0.00E+00 | 3185 | 866.5 |
| PA2458 | hypothetical protein | -5.44 | 1.70E-42 | 4.74E-41 | 366.5 | 96 |
| ohr | organic hydroperoxide resistance protein | -5.59 | 4.80E-24 | 9.27E-23 | 290 | 74 |
| PA1597 | hypothetical protein | -5.64 | 1.23E-19 | 1.97E-18 | 1500.5 | 381.5 |
| putP | sodium/proline symporter PutP | -5.67 | 0.00E+00 | 0.00E+00 | 7643 | 1939.5 |
| speC | ornithine decarboxylase | -5.69 | 0.00E+00 | 0.00E+00 | 2695 | 680.5 |
| phzC2 | phenazine biosynthesis protein PhzC | -5.69 | 4.27E-07 | 2.15E-06 | 38 | 9.5 |
| secB | secretion protein SecB | -5.70 | 0.00E+00 | 0.00E+00 | 4310.5 | 1085 |
| PA2916 | hypothetical protein | -5.70 | 1.56E-09 | 1.08E-08 | 48 | 12 |
| groEL | GroEL protein | -5.76 | 0.00E+00 | 0.00E+00 | 84401.5 | 21089 |
| rgsA | RgsA | -5.78 | 8.27E-29 | 1.83E-27 | 389 | 96.5 |
| PA2463 | hypothetical protein | -5.81 | 1.40E-12 | 1.29E-11 | 406.5 | 100 |
| PA4200 | hypothetical protein | -5.85 | 5.00E-72 | 1.83E-70 | 1475.5 | 361.5 |
| PA0505 | hypothetical protein | -5.88 | 1.97E-32 | 4.84E-31 | 393.5 | 95.5 |
| piv | protease IV | -5.89 | 0.00E+00 | 0.00E+00 | 4489 | 1095.5 |
| PA4128 | conserved hypothetical protein | -5.91 | 9.87E-77 | 3.93E-75 | 732 | 177.5 |
| tufA | elongation factor Tu | -5.97 | 0.00E+00 | 0.00E+00 | 57569 | 13871.5 |
| metK | methionine adenosyltransferase | -5.98 | 0.00E+00 | 0.00E+00 | 9020.5 | 2162.5 |
| PA0099 | type VI effector protein | -6.16 | 7.95E-21 | 1.35E-19 | 204 | 47.5 |
| PA1202 | probable hydrolase | -6.21 | 3.91E-28 | 8.54E-27 | 2382.5 | 550.5 |
| PA0496 | conserved hypothetical protein | -6.25 | 3.54E-17 | 4.96E-16 | 594 | 135.5 |
| PA2457 | hypothetical protein | -6.27 | 5.10E-18 | 7.44E-17 | 83.5 | 19 |
| PA4223 | probable ATP-binding component of ABC transporter | -6.28 | 3.73E-06 | 1.64E-05 | 804.5 | 178 |
| PA0603 | AgtA | -6.34 | 5.71E-04 | 1.63E-03 | 2213 | 493.5 |
| rpsR | 30S ribosomal protein S18 | -6.46 | 0.00E+00 | 0.00E+00 | 6160 | 1371 |
| PA0098 | hypothetical protein | -6.59 | 6.74E-18 | 9.77E-17 | 97 | 21 |
| pvdE | pyoverdine biosynthesis protein PvdE | -6.74 | 3.56E-16 | 4.61E-15 | 85 | 18 |
| PA0784 | probable transcriptional regulator | -6.77 | 2.30E-74 | 8.98E-73 | 1077.5 | 227 |
| speD | S-adenosylmethionine decarboxylase proenzyme | -6.92 | 3.24E-16 | 4.23E-15 | 1744 | 362 |
| oprC | Putative copper transport outer membrane porin OprC precursor | -6.92 | 0.00E+00 | 0.00E+00 | 4619 | 960 |
| PA0131 | BauB | -6.95 | 3.97E-32 | 9.72E-31 | 465 | 96 |
| PA0529 | conserved hypothetical protein | -6.98 | 2.58E-78 | 1.07E-76 | 2232.5 | 458.5 |
| rpsF | 30S ribosomal protein S6 | -7.17 | 0.00E+00 | 0.00E+00 | 19285 | 3874.5 |
| PA0050 | hypothetical protein | -7.31 | 2.72E-05 | 1.03E-04 | 13.5 | 2.5 |
| cioA | cyanide insensitive terminal oxidase | -7.39 | 1.42E-75 | 5.63E-74 | 1874.5 | 364 |
| PA4222 | probable ATP-binding component of ABC transporter | -7.46 | 8.98E-10 | 6.42E-09 | 798.5 | 149.5 |
| PA3961 | probable ATP-dependent helicase | -7.49 | 5.50E-29 | 1.23E-27 | 973.5 | 185.5 |
| fliC | flagellin type B | -7.77 | 0.00E+00 | 0.00E+00 | 41854.5 | 7728 |
| PA4277.3 | tRNA-Tyr | -7.84 | 1.98E-02 | 3.88E-02 | 3.5 | 0.5 |
| fptA | Fe(III)-pyochelin outer membrane receptor precursor | -7.98 | 3.46E-16 | 4.49E-15 | 1706.5 | 300.5 |
| flgL | flagellar hook-associated protein type 3 FlgL | -8.03 | 2.01E-74 | 7.91E-73 | 1321 | 236.5 |
| PA0495 | hypothetical protein | -8.09 | 9.87E-12 | 8.47E-11 | 553 | 97.5 |
| PA5318 | hypothetical protein | -8.42 | 3.59E-22 | 6.53E-21 | 150.5 | 25.5 |
| pchG | pyochelin biosynthetic protein PchG | -8.42 | 8.07E-08 | 4.60E-07 | 811 | 134 |
| hcpB | secreted protein Hcp | -8.53 | 1.29E-63 | 4.40E-62 | 1707.5 | 287 |
| PA2294 | probable ATP-binding component of ABC transporter | -8.78 | 6.03E-10 | 4.39E-09 | 28.5 | 4.5 |
| PA2462 | hypothetical protein | -8.80 | 0.00E+00 | 0.00E+00 | 7134.5 | 1159.5 |
| rsmY | regulatory RNA RsmY | -9.11 | 2.48E-05 | 9.47E-05 | 89 | 13.5 |
| PA2774 | Tse4 | -9.27 | 6.02E-08 | 3.48E-07 | 183 | 28 |
| pchF | pyochelin synthetase | -9.66 | 3.66E-10 | 2.76E-09 | 5641 | 814.5 |
| pchE | dihydroaeruginoic acid synthetase | -10.07 | 5.37E-11 | 4.39E-10 | 4239.5 | 588 |
| PA2136 | hypothetical protein | -10.15 | 4.41E-15 | 5.22E-14 | 72.5 | 10 |
| PA4514 | probable outer membrane receptor for iron transport | -10.35 | 9.61E-78 | 3.94E-76 | 522 | 72 |
| PA0130 | 3-Oxopropanoate dehydrogenase | -10.63 | 3.34E-59 | 1.10E-57 | 4412.5 | 595 |
| PA4220 | hypothetical protein | -10.91 | 2.33E-15 | 2.82E-14 | 66 | 8.5 |
| PA1244 | QslA, LasR-specific antiactivator QslA chain E | -11.40 | 9.63E-74 | 3.68E-72 | 611 | 77 |
| PA0494 | probable acyl-CoA carboxylase subunit | -11.62 | 7.53E-14 | 7.91E-13 | 773 | 94 |
| pncB1 | nicotinate phosphoribosyltransferase | -12.14 | 2.59E-101 | 1.29E-99 | 4381 | 516 |
| PA0498 | hypothetical protein | -13.49 | 2.96E-15 | 3.55E-14 | 48.5 | 5 |
| PA0493 | probable biotin-requiring enzyme | -14.39 | 2.84E-09 | 1.90E-08 | 123 | 12 |
| pvdJ | PvdJ | -14.95 | 3.53E-74 | 1.36E-72 | 466 | 44.5 |
| PA0132 | Beta-alanine:pyruvate transaminase | -15.25 | 6.18E-25 | 1.25E-23 | 4173.5 | 393 |
| PA3143 | transposase | -16.12 | 3.26E-25 | 6.67E-24 | 154.5 | 13.5 |
| PA0497 | hypothetical protein | -18.29 | 5.03E-16 | 6.43E-15 | 129.5 | 10 |
| waaL | O-antigen ligase, WaaL | -18.84 | 1.84E-18 | 2.74E-17 | 520.5 | 39.5 |
| PA2184 | conserved hypothetical protein | -20.41 | 1.50E-07 | 8.21E-07 | 16.5 | 1 |
| putA | proline dehydrogenase PutA | -20.61 | 0.00E+00 | 0.00E+00 | 19522 | 1362 |
| PA1508 | hypothetical protein | -22.67 | 6.57E-21 | 1.13E-19 | 65.5 | 4 |
| PA4134 | hypothetical protein | -24.57 | 6.13E-91 | 2.86E-89 | 516 | 30 |
| PA4129 | hypothetical protein | -25.40 | 1.14E-114 | 6.39E-113 | 2357 | 133 |
| pvdD | pyoverdine synthetase D | -25.66 | 3.15E-91 | 1.49E-89 | 468 | 26 |
| PA4132 | conserved hypothetical protein | -26.13 | 8.59E-134 | 5.32E-132 | 10256 | 561.5 |
| PA1316 | probable major facilitator superfamily (MFS) transporter | -26.32 | 2.59E-28 | 5.68E-27 | 76.5 | 4 |
| PA2427 | hypothetical protein | -30.84 | 2.67E-07 | 1.40E-06 | 14 | 0.5 |
| PA4130 | probable sulfite or nitrite reductase | -32.71 | 2.97E-141 | 1.88E-139 | 10002.5 | 437.5 |
| PA4918 | nicotinamidase, PcnA | -35.29 | 8.26E-146 | 5.54E-144 | 2787 | 113 |
| PA2159 | conserved hypothetical protein | -37.10 | 5.19E-03 | 1.19E-02 | 4 | 0 |
| PA5088 | type VI secretion lipase immunity protein, Tli5b3 | -38.86 | 1.99E-06 | 9.19E-06 | 181 | 6.5 |
| PA3939 | hypothetical protein | -42.32 | 1.35E-39 | 3.67E-38 | 108 | 3.5 |
| PA2158 | probable alcohol dehydrogenase (Zn-dependent) | -43.74 | 8.28E-11 | 6.62E-10 | 20 | 0.5 |
| PA2190 | conserved hypothetical protein | -46.95 | 7.78E-12 | 6.73E-11 | 21.5 | 0.5 |
| PA2335 | probable TonB-dependent receptor | -48.04 | 9.37E-12 | 8.05E-11 | 22 | 0.5 |
| PA0725 | hypothetical protein of bacteriophage Pf1 | -64.24 | 1.31E-04 | 4.31E-04 | 7 | 0 |
| PA2597 | hypothetical protein | -66.80 | 8.80E-24 | 1.68E-22 | 54 | 1 |
| fgtA | flagellar glycosyl transferase, FgtA | -67.33 | 5.84E-142 | 3.74E-140 | 4585.5 | 98 |
| PA1231 | conserved hypothetical protein | -68.71 | 7.19E-05 | 2.52E-04 | 7.5 | 0 |
| PA1196 | transcriptional regulator DdaR | -72.53 | 3.00E-132 | 1.84E-130 | 1147 | 22.5 |
| PA2295 | probable permease of ABC transporter | -73.16 | 4.25E-05 | 1.55E-04 | 8 | 0 |
| PA4133 | cytochrome c oxidase subunit (cbb3-type) | -73.31 | 5.15E-110 | 2.82E-108 | 8438 | 166 |
| PA2772a |  | -82.19 | 1.29E-05 | 5.19E-05 | 9 | 0 |
| PA2336 | hypothetical protein | -82.31 | 1.21E-05 | 4.91E-05 | 9 | 0 |
| PA2192 | conserved hypothetical protein | -91.44 | 2.86E-06 | 1.28E-05 | 10 | 0 |
| PA2333 | probable sulfatase | -92.53 | 5.40E-18 | 7.86E-17 | 42.5 | 0.5 |
| PA4131 | probable iron-sulfur protein | -93.47 | 5.69E-187 | 5.41E-185 | 13653.5 | 209 |
| PA2182 | hypothetical protein | -100.24 | 7.53E-07 | 3.68E-06 | 11 | 0 |
| PA1233 | hypothetical protein | -105.69 | 3.98E-23 | 7.43E-22 | 48.5 | 0.5 |
| PA3434 | probable transposase | -114.27 | 5.12E-07 | 2.55E-06 | 12.5 | 0 |
| PA2334 | probable transcriptional regulator | -122.60 | 1.51E-07 | 8.26E-07 | 13.5 | 0 |
| PA2188 | probable alcohol dehydrogenase (Zn-dependent) | -131.80 | 1.91E-08 | 1.17E-07 | 14.5 | 0 |
| PA2108 | probable decarboxylase | -155.84 | 4.60E-29 | 1.04E-27 | 71.5 | 0.5 |
| PA2183 | hypothetical protein | -163.42 | 2.80E-10 | 2.13E-09 | 18 | 0 |
| PA1313 | probable major facilitator superfamily (MFS) transporter | -163.84 | 8.47E-10 | 6.09E-09 | 18 | 0 |
| PA3142 | integrase | -172.22 | 2.30E-55 | 7.28E-54 | 200 | 1.5 |
| PA0258 | hypothetical protein | -177.34 | 1.32E-10 | 1.04E-09 | 19.5 | 0 |
| PA0457.1 | hypothetical membrane protein | -179.06 | 9.38E-30 | 2.15E-28 | 82.5 | 0.5 |
| PA1931 | probable ferredoxin | -186.72 | 2.54E-10 | 1.94E-09 | 20.5 | 0 |
| PA2602 | 3-mercaptopropionate dioxygenase | -190.70 | 1.31E-11 | 1.12E-10 | 21 | 0 |
| PA3869 | hypothetical protein | -190.77 | 1.65E-11 | 1.39E-10 | 21 | 0 |
| PA4148 | probable short-chain dehydrogenase | -195.34 | 2.13E-11 | 1.78E-10 | 21.5 | 0 |
| PA4152 | probable hydrolase | -198.48 | 1.44E-09 | 1.00E-08 | 22 | 0 |
| PA1314 | hypothetical protein | -199.67 | 4.80E-12 | 4.24E-11 | 22 | 0 |
| PA0718 | hypothetical protein of bacteriophage Pf1 | -204.29 | 2.05E-12 | 1.86E-11 | 22.5 | 0 |
| PA1472 | conserved hypothetical protein | -213.52 | 2.25E-39 | 6.07E-38 | 98 | 0.5 |
| PA1239 | hypothetical protein | -217.66 | 5.68E-13 | 5.38E-12 | 24 | 0 |
| PA2565 | hypothetical protein | -218.52 | 1.99E-11 | 1.67E-10 | 24 | 0 |
| PA3065 | hypothetical protein | -235.87 | 6.74E-14 | 7.12E-13 | 26 | 0 |
| katN | non-heme catalase KatN | -236.27 | 1.68E-12 | 1.54E-11 | 26 | 0 |
| PA0984 | colicin immunity protein | -245.26 | 1.39E-13 | 1.42E-12 | 27 | 0 |
| arr | aminoglycoside response regulator, inner membrane phosphodiesterase | -249.99 | 1.32E-55 | 4.20E-54 | 202.5 | 1 |
| PA1195 | dimethylarginine dimethylaminohydrolase DdaH | -260.54 | 6.67E-41 | 1.85E-39 | 119.5 | 0.5 |
| PA0204 | probable permease of ABC transporter | -261.87 | 3.93E-13 | 3.81E-12 | 29 | 0 |
| PA2073 | probable transporter (membrane subunit) | -267.71 | 3.31E-12 | 2.97E-11 | 29.5 | 0 |
| PA2109 | hypothetical protein | -268.63 | 3.49E-44 | 9.94E-43 | 123.5 | 0.5 |
| PA2461 | hypothetical protein | -279.03 | 9.73E-41 | 2.69E-39 | 128 | 0.5 |
| PA2596 | conserved hypothetical protein | -284.75 | 8.20E-15 | 9.45E-14 | 31.5 | 0 |
| PA3868 | hypothetical protein | -285.50 | 2.54E-16 | 3.34E-15 | 31.5 | 0 |
| PA4153 | 2,3-butanediol dehydrogenase | -285.60 | 8.67E-16 | 1.10E-14 | 31.5 | 0 |
| PA4194 | probable permease of ABC transporter | -315.99 | 5.79E-15 | 6.80E-14 | 35 | 0 |
| PA4193 | probable permease of ABC transporter | -316.31 | 1.08E-14 | 1.23E-13 | 35 | 0 |
| PA0206 | probable ATP-binding component of ABC transporter | -325.56 | 1.97E-17 | 2.79E-16 | 36 | 0 |
| PA0188 | hypothetical protein | -331.10 | 2.59E-16 | 3.40E-15 | 36.5 | 0 |
| PA0261 | Type 6 lipase immunity, Tli3 | -334.25 | 2.39E-24 | 4.69E-23 | 153 | 0.5 |
| PA2296 | hypothetical protein | -352.35 | 2.34E-14 | 2.60E-13 | 39 | 0 |
| acoB | acetoin catabolism protein AcoB | -357.17 | 4.36E-14 | 4.70E-13 | 39.5 | 0 |
| PA0203 | probable binding protein component of ABC transporter | -358.63 | 1.75E-17 | 2.48E-16 | 39.5 | 0 |
| fpvA | ferripyoverdine receptor | -358.96 | 7.66E-14 | 8.03E-13 | 2056 | 8 |
| PA0205 | probable permease of ABC transporter | -368.10 | 1.69E-16 | 2.28E-15 | 40.5 | 0 |
| PA1240 | probable enoyl-CoA hydratase/isomerase | -375.64 | 5.21E-20 | 8.53E-19 | 41.5 | 0 |
| treA | periplasmic trehalase precursor | -388.06 | 2.83E-17 | 3.98E-16 | 43 | 0 |
| PA0719 | hypothetical protein of bacteriophage Pf1 | -398.64 | 4.81E-21 | 8.38E-20 | 44 | 0 |
| PA2459 | hypothetical protein | -407.58 | 3.23E-12 | 2.90E-11 | 45 | 0 |
| exoY | adenylate cyclase ExoY | -415.56 | 1.26E-19 | 2.02E-18 | 46 | 0 |
| PA4192 | probable ATP-binding component of ABC transporter | -420.16 | 1.83E-20 | 3.04E-19 | 46.5 | 0 |
| PA1366 | hypothetical protein | -428.59 | 2.04E-89 | 9.24E-88 | 494.5 | 1.5 |
| PA2594 | conserved hypothetical protein | -432.30 | 5.02E-13 | 4.83E-12 | 199.5 | 0.5 |
| PA3067 | probable transcriptional regulator | -447.80 | 1.44E-23 | 2.73E-22 | 49.5 | 0 |
| PA0689 | low-molecular-weight alkaline phosphatase B, LapB | -476.04 | 1.30E-11 | 1.10E-10 | 218 | 0.5 |
| PA1932 | probable hydroxylase molybdopterin-containing subunit | -479.45 | 3.94E-24 | 7.63E-23 | 53 | 0 |
| PA1095 | hypothetical protein | -482.71 | 4.29E-178 | 3.54E-176 | 3077 | 9 |
| PA4150 | probable dehydrogenase E1 component | -487.93 | 4.31E-19 | 6.72E-18 | 54 | 0 |
| PA0715 | hypothetical protein | -519.02 | 7.98E-137 | 5.00E-135 | 781 | 2 |
| coaB | coat protein B of bacteriophage Pf1 | -534.29 | 1.72E-97 | 8.46E-96 | 431.5 | 1 |
| PA4149 | conserved hypothetical protein | -541.97 | 1.28E-24 | 2.56E-23 | 60 | 0 |
| PA2595 | conserved hypothetical protein | -542.77 | 1.82E-17 | 2.58E-16 | 60 | 0 |
| PA3867 | probable DNA invertase | -543.39 | 2.43E-26 | 5.12E-25 | 60 | 0 |
| PA0724 | probable coat protein A of bacteriophage Pf1 | -551.70 | 6.15E-28 | 1.33E-26 | 61 | 0 |
| PA2102 | hypothetical protein | -553.08 | 1.58E-24 | 3.13E-23 | 61 | 0 |
| PA2103 | probable molybdopterin biosynthesis protein MoeB | -569.66 | 2.24E-70 | 7.97E-69 | 261.5 | 0.5 |
| PA2775 | Tsi4 | -601.40 | 1.05E-08 | 6.63E-08 | 66.5 | 0 |
| PA2599 | conserved hypothetical protein | -612.34 | 8.01E-22 | 1.44E-20 | 67.5 | 0 |
| PA1232 | hypothetical protein | -619.82 | 1.26E-27 | 2.70E-26 | 68.5 | 0 |
| PA4195 | probable binding protein component of ABC transporter | -642.53 | 3.47E-22 | 6.34E-21 | 71 | 0 |
| PA2600 | hypothetical protein | -652.85 | 1.07E-24 | 2.15E-23 | 72 | 0 |
| PA0187 | hypothetical protein | -654.93 | 3.66E-30 | 8.47E-29 | 72.5 | 0 |
| PA1471 | hypothetical protein | -659.15 | 1.05E-29 | 2.41E-28 | 73 | 0 |
| PA4849 | hypothetical protein | -667.89 | 9.82E-08 | 5.51E-07 | 74 | 0 |
| PA2772 | hypothetical protein | -668.75 | 2.64E-30 | 6.14E-29 | 74 | 0 |
| PA1229 | probable transcriptional regulator | -681.04 | 1.53E-23 | 2.88E-22 | 75.5 | 0 |
| PA1088 | hypothetical protein | -684.65 | 1.94E-131 | 1.16E-129 | 554 | 1 |
| PA2771 | diguanylate cyclase with a self-blocked I-site, Dcsbis | -690.82 | 3.77E-16 | 4.87E-15 | 318.5 | 0.5 |
| PA4802 | hypothetical protein | -696.32 | 1.69E-16 | 2.28E-15 | 77 | 0 |
| acoR | transcriptional regulator AcoR | -703.31 | 4.83E-15 | 5.71E-14 | 324.5 | 0.5 |
| pys2 | pyocin S2 | -711.12 | 1.15E-13 | 1.18E-12 | 1809 | 3.5 |
| PA0722 | hypothetical protein of bacteriophage Pf1 | -746.60 | 2.03E-33 | 5.14E-32 | 82.5 | 0 |
| PA3865a |  | -767.36 | 4.56E-33 | 1.14E-31 | 85 | 0 |
| PA2598 | hypothetical protein | -792.28 | 1.91E-33 | 4.87E-32 | 87.5 | 0 |
| PA3066 | hypothetical protein | -792.70 | 4.23E-32 | 1.03E-30 | 87.5 | 0 |
| PA0726 | hypothetical protein of bacteriophage Pf1 | -801.89 | 1.23E-31 | 2.97E-30 | 88.5 | 0 |
| PA2566 | conserved hypothetical protein | -859.60 | 1.08E-35 | 2.81E-34 | 95 | 0 |
| PA0716 | hypothetical protein | -895.22 | 1.23E-119 | 7.10E-118 | 723 | 1 |
| PA1315 | probable transcriptional regulator | -908.33 | 8.73E-15 | 1.00E-13 | 100.5 | 0 |
| PA1312 | probable transcriptional regulator | -908.64 | 2.69E-23 | 5.06E-22 | 100.5 | 0 |
| PA2564 | hypothetical protein | -916.93 | 1.97E-26 | 4.16E-25 | 101.5 | 0 |
| PA1194 | probable amino acid permease | -923.36 | 1.13E-17 | 1.62E-16 | 102.5 | 0 |
| imm2 | pyocin S2 immunity protein | -926.25 | 1.72E-96 | 8.28E-95 | 747 | 1 |
| PA1241 | probable transcriptional regulator | -1000.88 | 1.04E-07 | 5.82E-07 | 111 | 0 |
| PA0701a |  | -1021.02 | 1.41E-29 | 3.22E-28 | 113 | 0 |
| wbpL | glycosyltransferase WbpL | -1048.06 | 1.72E-177 | 1.40E-175 | 1212.5 | 1.5 |
| PA2460 | hypothetical protein | -1057.01 | 9.64E-33 | 2.39E-31 | 116.5 | 0 |
| PA1371 | hypothetical protein | -1074.07 | 2.12E-77 | 8.50E-76 | 491.5 | 0.5 |
| pilA | type 4 fimbrial precursor PilA | -1086.08 | 2.50E-231 | 2.64E-229 | 36909.5 | 48.5 |
| PA0260 | Type 6 lipase effector, Tle3 | -1104.67 | 1.05E-14 | 1.20E-13 | 1272.5 | 1.5 |
| PA1368 | hypothetical protein | -1175.21 | 1.38E-49 | 4.18E-48 | 130 | 0 |
| PA2106 | hypothetical protein | -1197.61 | 1.14E-50 | 3.52E-49 | 132.5 | 0 |
| PA2037 | hypothetical protein | -1206.53 | 1.64E-10 | 1.28E-09 | 133 | 0 |
| wbpH | probable glycosyltransferase WbpH | -1239.27 | 9.90E-179 | 8.29E-177 | 2296.5 | 2.5 |
| PA1372 | hypothetical protein | -1258.07 | 6.74E-118 | 3.80E-116 | 3189 | 3.5 |
| PA0257 | hypothetical protein | -1267.06 | 2.17E-09 | 1.47E-08 | 140 | 0 |
| PA0634 | hypothetical protein | -1304.32 | 1.07E-145 | 7.06E-144 | 1053 | 1 |
| wbpJ | probable glycosyl transferase WbpJ | -1334.51 | 1.05E-174 | 8.19E-173 | 1543.5 | 1.5 |
| PA1093 | hypothetical protein | -1409.42 | 5.04E-185 | 4.63E-183 | 2120.5 | 2 |
| PA0645 | hypothetical protein | -1441.71 | 3.23E-58 | 1.05E-56 | 159.5 | 0 |
| PA0640 | probable bacteriophage protein | -1450.51 | 4.86E-112 | 2.69E-110 | 665 | 0.5 |
| PA0639 | conserved hypothetical protein | -1472.67 | 6.52E-131 | 3.87E-129 | 676 | 0.5 |
| PA2105 | probable acetyltransferase | -1511.02 | 5.09E-55 | 1.59E-53 | 167 | 0 |
| hisF2 | imidazoleglycerol-phosphate synthase, cyclase subunit | -1535.70 | 2.35E-156 | 1.72E-154 | 1240 | 1 |
| PA2104 | probable cysteine synthase | -1592.76 | 6.12E-56 | 1.96E-54 | 176 | 0 |
| PA4191 | isopenicillin-N synthase | -1605.13 | 5.69E-23 | 1.06E-21 | 177.5 | 0 |
| PA2734 | hypothetical protein | -1607.03 | 5.57E-185 | 5.04E-183 | 1859.5 | 1.5 |
| PA2736 | hypothetical protein | -1611.53 | 8.00E-62 | 2.71E-60 | 178.5 | 0 |
| PA0648 | hypothetical protein | -1613.99 | 1.69E-61 | 5.68E-60 | 178.5 | 0 |
| PA0646 | hypothetical protein | -1627.40 | 4.55E-144 | 2.95E-142 | 1313 | 1 |
| PA2764 | hypothetical protein | -1702.48 | 4.34E-13 | 4.19E-12 | 188 | 0 |
| PA2601 | probable transcriptional regulator | -1703.81 | 3.54E-23 | 6.64E-22 | 189 | 0 |
| PA1370 | hypothetical protein | -1762.13 | 2.94E-102 | 1.50E-100 | 807.5 | 0.5 |
| PA2603 | probable thiosulfate sulfurtransferase | -1773.64 | 3.80E-50 | 1.16E-48 | 196.5 | 0 |
| PA2730 | hypothetical protein | -1778.55 | 1.34E-146 | 9.07E-145 | 817 | 0.5 |
| PA1939 | hypothetical protein | -1901.99 | 4.74E-161 | 3.55E-159 | 3519 | 2.5 |
| PA3362 | hypothetical protein | -1911.94 | 5.35E-51 | 1.66E-49 | 211.5 | 0 |
| fliD | flagellar capping protein FliD | -1940.68 | 8.78E-223 | 9.09E-221 | 6996 | 5 |
| PA0259 | Type 6 lipase adaptor, Tla3 | -1965.10 | 8.94E-12 | 7.69E-11 | 908.5 | 0.5 |
| PA3157 | probable acetyltransferase | -2002.13 | 1.79E-155 | 1.29E-153 | 921.5 | 0.5 |
| PA1933 | probable hydroxylase large subunit | -2039.05 | 1.74E-147 | 1.19E-145 | 939 | 0.5 |
| PA0720 | helix destabilizing protein of bacteriophage Pf1 | -2061.32 | 4.69E-72 | 1.72E-70 | 228 | 0 |
| PA1428a |  | -2074.17 | 2.82E-24 | 5.52E-23 | 229.5 | 0 |
| wbpE | UDP-2-acetamido-2-dideoxy-d-ribo-hex-3-uluronic acid transaminase, wbpE | -2162.38 | 4.00E-185 | 3.73E-183 | 4751.5 | 3 |
| wbpB | UDP-2-acetamido-2-deoxy-d-glucuronic acid 3-dehydrogenase, WbpB | -2179.93 | 2.11E-176 | 1.70E-174 | 4038.5 | 2.5 |
| wbpG | LPS biosynthesis protein WbpG | -2295.22 | 1.54E-151 | 1.08E-149 | 4243 | 2.5 |
| PA2732 | hypothetical protein | -2320.92 | 7.83E-188 | 7.56E-186 | 2685 | 1.5 |
| wbpA | UDP-N-acetyl-d-glucosamine 6-Dehydrogenase | -2360.08 | 1.36E-216 | 1.39E-214 | 6851 | 4 |
| wzz | O-antigen chain length regulator | -2400.52 | 3.08E-166 | 2.37E-164 | 1103 | 0.5 |
| PA0207 | probable transcriptional regulator | -2499.05 | 2.89E-83 | 1.25E-81 | 276.5 | 0 |
| PA1090 | hypothetical protein | -2538.24 | 1.43E-85 | 6.26E-84 | 281 | 0 |
| PA0647 | hypothetical protein | -2565.71 | 3.30E-60 | 1.10E-58 | 283 | 0 |
| PA0561 | hypothetical protein | -2655.50 | 8.46E-14 | 8.78E-13 | 294 | 0 |
| PA0202 | probable amidase | -2687.17 | 3.81E-64 | 1.31E-62 | 298.5 | 0 |
| PA5265 | hypothetical protein | -2708.91 | 7.13E-13 | 6.72E-12 | 1250.5 | 0.5 |
| PA0053 | hypothetical protein | -2988.50 | 1.27E-21 | 2.23E-20 | 330.5 | 0 |
| PA1089 | conserved hypothetical protein | -3066.03 | 1.82E-89 | 8.30E-88 | 339 | 0 |
| PA3866 | Pyocin S4 | -3066.66 | 5.60E-184 | 4.98E-182 | 3549.5 | 1.5 |
| exoS | exoenzyme S | -3316.73 | 1.20E-77 | 4.90E-76 | 366.5 | 0 |
| PA1369 | hypothetical protein | -3317.01 | 2.77E-74 | 1.07E-72 | 366 | 0 |
| PA1096 | hypothetical protein | -3395.06 | 6.22E-179 | 5.29E-177 | 1560.5 | 0.5 |
| PA0644 | hypothetical protein | -3414.45 | 2.26E-97 | 1.10E-95 | 377.5 | 0 |
| PA2733 | conserved hypothetical protein | -3493.84 | 1.27E-90 | 5.86E-89 | 386 | 0 |
| PA0637 | conserved hypothetical protein | -3566.94 | 7.44E-104 | 3.86E-102 | 394.5 | 0 |
| wzx | O-antigen translocase | -3664.63 | 1.74E-109 | 9.43E-108 | 406 | 0 |
| PA3488 | Tli5 | -3889.09 | 3.84E-78 | 1.58E-76 | 429 | 0 |
| hisH2 | glutamine amidotransferase | -4086.10 | 6.30E-87 | 2.80E-85 | 451 | 0 |
| PA0100 | hypothetical protein | -4197.29 | 5.74E-108 | 3.08E-106 | 464 | 0 |
| PA5264 | hypothetical protein | -4331.88 | 8.04E-14 | 8.39E-13 | 478.5 | 0 |
| PA0641 | probable bacteriophage protein | -4606.15 | 4.09E-198 | 4.09E-196 | 3727.5 | 1 |
| PA0635 | hypothetical protein | -5248.32 | 4.24E-127 | 2.49E-125 | 580.5 | 0 |
| PA0820 | hypothetical protein | -5333.95 | 2.74E-105 | 1.45E-103 | 589 | 0 |
| PA2119 | alcohol dehydrogenase (Zn-dependent) | -5835.59 | 4.30E-15 | 5.10E-14 | 4669.5 | 1 |
| PA0638 | probable bacteriophage protein | -6198.24 | 1.56E-144 | 1.02E-142 | 686 | 0 |
| PA2735 | type I HsdM methyltransferase | -6365.31 | 1.13E-191 | 1.11E-189 | 2928 | 0.5 |
| wzy | B-band O-antigen polymerase | -6615.39 | 3.66E-150 | 2.54E-148 | 732.5 | 0 |
| wbpI | UDP-N-acetylglucosamine 2-epimerase WbpI | -7555.15 | 1.56E-157 | 1.15E-155 | 3462 | 0.5 |
| PA0643 | hypothetical protein | -9568.88 | 6.00E-132 | 3.63E-130 | 1056.5 | 0 |
| wbpK | probable NAD-dependent epimerase/dehydratase WbpK | -10309.97 | 2.10E-162 | 1.60E-160 | 1140 | 0 |
| pyoS5 | pyocin S5 | -13501.20 | 7.54E-127 | 4.38E-125 | 1489.5 | 0 |
| wbpD | UDP-2-acetamido-3-amino-2,3-dideoxy-d-glucuronic acid N-acetyltransferase, WbpD | -15416.88 | 8.61E-176 | 6.81E-174 | 1705.5 | 0 |
| PA0633 | hypothetical protein | -20240.88 | 3.49E-180 | 3.06E-178 | 2238 | 0 |
| pldA | lipase effector of T6SS | -22643.87 | 6.25E-152 | 4.45E-150 | 2500 | 0 |
| PA0636 | hypothetical protein | -25987.93 | 1.47E-179 | 1.27E-177 | 2873.5 | 0 |

**Table S7. Significantly regulated genes in Iso 2 comparing to that of PAO1 based on the selection criteria of absolute change≧4 and adjusted p-value<0.05.**
