## Supplementary Table S8 for "Persistent bacterial coinfection of a COVID-19 patient caused by a genetically adapted *Pseudomonas aeruginosa* chronic colonizer"

|  | Iso 1 | Iso 2 |
| --- | --- | --- |
| ceftazidime-MIC | 19 S | <=1 S |
| ceftazidime-KB | 25mm S | 25mm S |
| piperacillin-MIC | 27 S | <=4 S |
| piperacillin-KB | 26mm S | 26mm S |
| cefoperazone/sulbactam-KB | 22mm S | 24mm S |
| imipenem-MIC | 32 S | 4 I |
| imipenem-KB | 27mm S | 9mm R |
| aztreonam-MIC | 22 S | <=2 S |
| aztreonam-KB | 22mm S | 21mm I |
| levofloxacin-MIC | 25 S | <=1 S |
| levofloxacin-KB | 21mm I | 19mm I |

**Table S8.** Results of antimicrobial susceptibility tests of the isolates.
