## Supplementray figures for "Persistent bacterial coinfection of a COVID-19 patient caused by a genetically adapted *Pseudomonas aeruginosa* chronic colonizer"

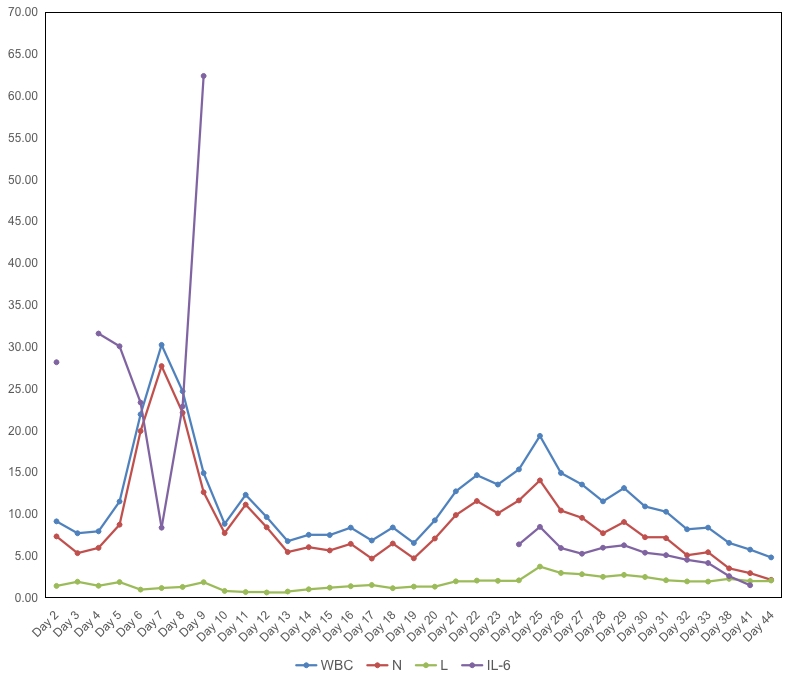

**Figure S1**.Levels of white blood cell, neutrophil, lymphocyte and interleukin-6 during the course of hospitalization.

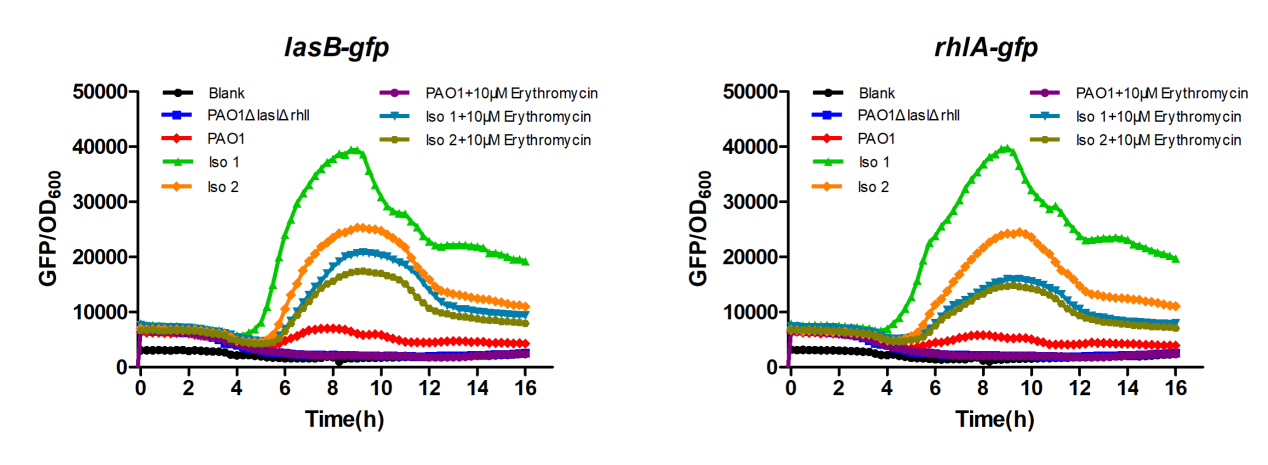

**Figure S1**.Inhibition of *las* and *rhl* quorum sensing by erythromycin.
